# DoFormer: Causal Transformer for Gene Perturbation

**DOI:** 10.64898/2026.05.02.722054

**Authors:** Alireza Karbalayghareh, Evan Paull, Andrea Califano

**Affiliations:** Biohub, New York, NY; Biohub & Columbia University, New York, NY

## Abstract

Learning causal gene regulatory mechanisms from single-cell data, and thereby predicting the effects of unseen perturbations, remains challenging. Observational RNA-seq data alone is insufficient for causal modeling, whereas perturbational data is essential. Classical causal inference methods often rely on unrealistic directed acyclic graph (DAG) assumptions and are not well suited to integrating multimodal data. Current transcriptomic foundation models also typically treat observational and perturbational data identically, limiting their ability to model perturbations. We present **DoFormer**, a causal multimodal Transformer that makes no DAG assumptions and leverages rich perturbational data to accurately predict previously unseen perturbations. DoFormer enables principled *in silico* perturbations by adapting the causal *do*-operator within the attention mechanism: the perturbed gene is set to the intervention value and prevented from attending to other genes, allowing the model to fully distinguish observational from interventional regimes. We train DoFormer using biologically informed loss functions and evaluate it with comprehensive perturbation prediction metrics. DoFormer substantially improves perturbation prediction relative to baseline and prior foundation models, underscoring the importance of intervention-aware architectures and biologically grounded objectives for causal modeling in single-cell genomics.

## 1 Introduction

Advances in scalable, fast Transformer-based AI models (Vaswani et al. [2017], Brown et al. [2020], Touvron et al. [2023], Dao [2023], Shah et al. [2024]) and the emergence of atlas-level single-cell RNA expression data across many cell types and tissues (Program et al. [2025], Replogle et al. [2022], Nadig et al. [2025], Zhang et al. [2025], Youngblut et al. [2025]) have led to the development of numerous transcriptomic foundation models aimed at accurately predicting out-of-context as well as unseen cellular perturbations. A central question for these AI models is whether they can advance our understanding of the causal biological mechanisms of cells in health and disease. Before the era of foundation models, causal discovery and inference provided a reliable framework for mechanistic understanding with great promise under restrictive assumptions (Pearl et al. [2000], Spirtes et al. [2000], Peters et al. [2017]). However, due to the probabilistic nature of the framework, classical causal discovery methods have two main limitations. First, they often assume a directed acyclic graph (DAG), which requires highly unrealistic assumptions about gene regulatory networks (GRNs), which frequently contain cycles and feedback loops. Second, they cannot incorporate multiple data modalities, greatly limiting the use of increasingly abundant and valuable datasets that allow us to view different aspects of the cellular phenotype simultaneously–a potentially revolutionary set of technologies.

Here, we present DoFormer, a causal multi-modal AI model without any DAG assumption, which uses two complementary sources of information for each gene: RNA expression and the sequence information of its protein, via protein language model (PLM) embeddings. We believe the following four features are required for any reliable causal AI model: 1) scalability, 2) multi-modality, 3) biologically focused loss functions, and 4) correct *in silico* perturbations. Thanks to Transformer-based architectures, current foundation models already scale and can accept PLM embeddings for genes. However, their fundamental limitation is the lack of proper *in silico* perturbation modeling. The main contribution of DoFormer is to show how to correctly model genetic perturbations.

We know from the causality literature that observational data alone are not sufficient to learn causal models Pearl et al. [2000]. Fortunately, we have the opportunity to leverage large-scale perturbational data (CRISPRi/a, KO, overexpression, etc.) in combination with observational scRNA-seq data to simultaneously learn gene–gene interactions and model *in silico* perturbations. Because the Transformer architecture operates at the gene level, it allows us to perform *in silico* perturbations using the *do*-operator, which we borrow from the causality literature to model *in silico* perturbations. This setup allows us to avoid making unreasonable assumptions that the underlying GRN is a DAG, and by training the model on the effects of perturbations, it will emphasize causal interactions and unlearn correlative interactions induced by scRNA-seq data alone.

Thanks to the causal inference literature, we have a mathematically grounded way of performing perturbations: the *do*-operator. In essence, if we intervene on a gene in a graph, we should remove all edges coming into the targeted gene and set its expression value to the intervened value. We adapted the *do*-operator in the Transformer architecture and consequently named the model DoFormer. We model *in silico* perturbations in DoFormer by not letting the perturbed gene attend to other genes and by setting its expression value to zero. In other words, the attention scores from the perturbed gene to all other genes are set to zero, allowing the model to distinguish whether the data are observational or perturbational and modify its structure accordingly. Existing foundation models have been trained in the same way on observational and perturbational data, and we believe that this is why they cannot outperform very simple baseline models (Ahlmann-Eltze et al. [2025], Kernfeld et al. [2025], Csendes et al. [2025]).

Another contribution of DoFormer is designing biologically focused loss functions based on the statistics of differential expression (DE) analyses of perturbations. One important note is that in perturbational data, we have perturbations in unmatched cells, whereas in the model we predict perturbations in the same cells. As a result, we should perform unpaired (two-sample) DE tests in real perturbations and paired (one-sample) DE tests in predicted perturbations. We split our training into two stages, pre-training and fine-tuning, as the model learns different tasks at each stage. We first pretrain DoFormer on observational control cells via masked language modeling (MLM) and then fine-tune it on perturbed cells using a weighted mean squared error (WMSE) loss, where weights come from the z-scores of the Wilcoxon rank-sum DE tests. In pretraining, the model learns general gene–gene interactions by predicting the RNA expression values of masked genes. These interactions are not necessarily causal, but they provide a meaningful initialization for fine-tuning. In fine-tuning, the model computes the *in silico* effect sizes and tries to match the true effect sizes, thereby refining the interactions toward causal ones.

## 2 Related work

Perturbation prediction has long been a goal of transcriptomic AI models. Models such as GEARS (Roohani et al. [2024]) build graph neural networks (GNNs) using knowledge of GRNs and Gene Ontology (GO) and suggest that these priors help improve perturbation prediction. Due to the success of large language models (LLMs), in recent years, we have seen several single-cell foundation models trained on large-scale observational RNA expression data (Cui et al. [2024], Theodoris et al. [2023], Hao et al. [2024], Yang et al. [2022]). These Transformer-based models are trained with MLM loss functions to reconstruct the RNA expression values of masked genes. The main hope for these foundation models is to learn rich representations of genes and cells in atlas-level observational datasets for various downstream tasks, such as cell type annotation and perturbation prediction. However, benchmarking these models has revealed that simple baselines, such as the mean of perturbed cells or a linear model, can sometimes outperform foundation models (Ahlmann-Eltze et al. [2025], Kernfeld et al. [2025], Csendes et al. [2025]).

A recent model, STATE (Adduri et al. [2025]), addresses some of the shortcomings of previous models by explicitly modeling perturbations in sets of control cells. STATE is a Transformer-based generative model that takes the RNA expression values of highly variable genes in a set of unperturbed control cells (in any cell type), maps them to a latent space, adds a perturbation token, and decodes the output of the Transformer block back to gene space to generate a set of perturbed cells using the MMD loss function. It performs attention across cells within a set and shows improved performance in perturbation prediction compared to other foundation models. One useful contribution of STATE is CELL-Eval, a comprehensive evaluation framework that tests a model’s ability to understand different biological metrics for any perturbation. We adapt CELL-Eval to our setting and show that DoFormer outperforms STATE and baseline models across these metrics. Since STATE outperforms other AI models, such as scGPT (Cui et al. [2024]), scVI (Lopez et al. [2018]), and CPA (Lotfollahi et al. [2023]), on the same dataset (Adduri et al. [2025]), we compare only against STATE and the baseline models (which often outperform STATE).

## 3 Methodology

### 3.1 Problem Formulation

Let 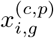 denote the log1p-normalized RNA expression value of gene *g* ∈ {1, …, *G*} in the cell *i* ∈ {1, …, *N*^(*c,p*)^} of the cell type *c* ∈ {1, …, *C*} under perturbation *p* ∈ {*ϕ*, 1, …, *P*}, where *p* = *ϕ* denotes the unperturbed observational state, and *N*^(*c,p*)^ is the number of cells in the cell type *c* under perturbation *p*. For simplicity, we drop the cell index *i* throughout the paper whenever possible and use 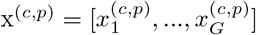 to represent a cell. The biological question that we are trying to answer is as follows: if we have access to a subset of cell types and perturbations, how well can we predict the effects of unseen perturbations in any cell type. Mathematically, if the model gets the control cells x^(*c,ϕ*)^ in the input, how well can it predict the effect sizes of any unseen perturbation 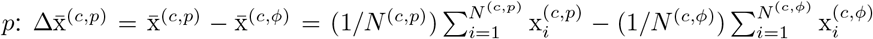. Note that, unlike many other methods that model perturbations by tokens, we do not assume that the model should see all the perturbations in at least one cell type. This makes DoFormer a very powerful model for predicting the effect sizes of all genetic perturbations in all cell types.

If we can build such a model that can predict accurate outcomes of any unseen perturbations, it would be of great use to biologists in designing experiments. Since the search space in biology is infinite, a causal transcriptomic AI model can tremendously shrink the space by nominating highly effective perturbation targets for lab-in-the-loop biological discoveries.

### 3.2 *Do*-operator in DoFormer

Transformers are universal function classes that have shown great performance in many tasks thanks to their global attention and scalability on HPC systems, and tools like flash attention (Dao [2023], Shah et al. [2024]) have made them a de-facto AI model class. Here we use Transformers and modify it by implementing the *do*-operator in the attention blocks to do the *in silico* perturbation. The *do*-operator can only be implemented in models that work in the gene space like Transformers, and the latent space models are not well-suited for it since they lose the gene resolution.

We have two states: observational *S*^(*ϕ*)^ and perturbational *S*^(*p*)^. The observational state does not have any *do*-operator, and the model is exactly the same as the Transformer. In the perturbational state *S*^(*p*)^, the model employs the *do*-operator to zero out the attention scores of the perturbed gene *p* to all other genes and to set its expression value in the input control cells to zero:

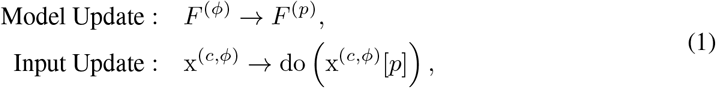

where do 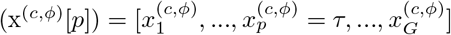, where *τ* is the RNA expression value of the gene *p* after perturbation. In the CRISPRi experiments, we use the default value of *τ* = 0, assuming complete knockout. In general, to make DoFormer suitable for any form of perturbational data (such as CRISPRi/a, overexpression, etc.), we define 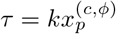, where *k* is the knockdown ratio (*k <* 1) or the activation ratio (*k >* 1).

*F*^(*ϕ*)^ is the original Transformer and *F*^(*p*)^ is the Transformer modified by the *do*-operator. Let 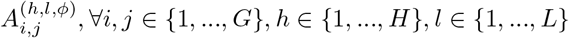, ∀*i, j* ∈ {1, …, *G*}, *h* ∈ {1, …, *H*}, *l* ∈ {1, …, *L*} denote the attention scores from the gene *i* to the gene *j* in the head *h* and the layer *l* in the observational model *F*^(*ϕ*)^. The attention scores in *F*^(*p*)^ with the input do(x^(*c,ϕ*)^[*p*]) are defined as:

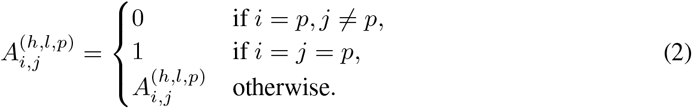

This is equivalent to masking the *p*-th row of the attention matrices to zeros except the *p*-th column. A very simplified schematic of the DoFormer’s attention blocks for a three-gene scenario is depicted in Fig. 1. The complete DoFormer architecture is shown in Fig. 29.

**Figure 1:**
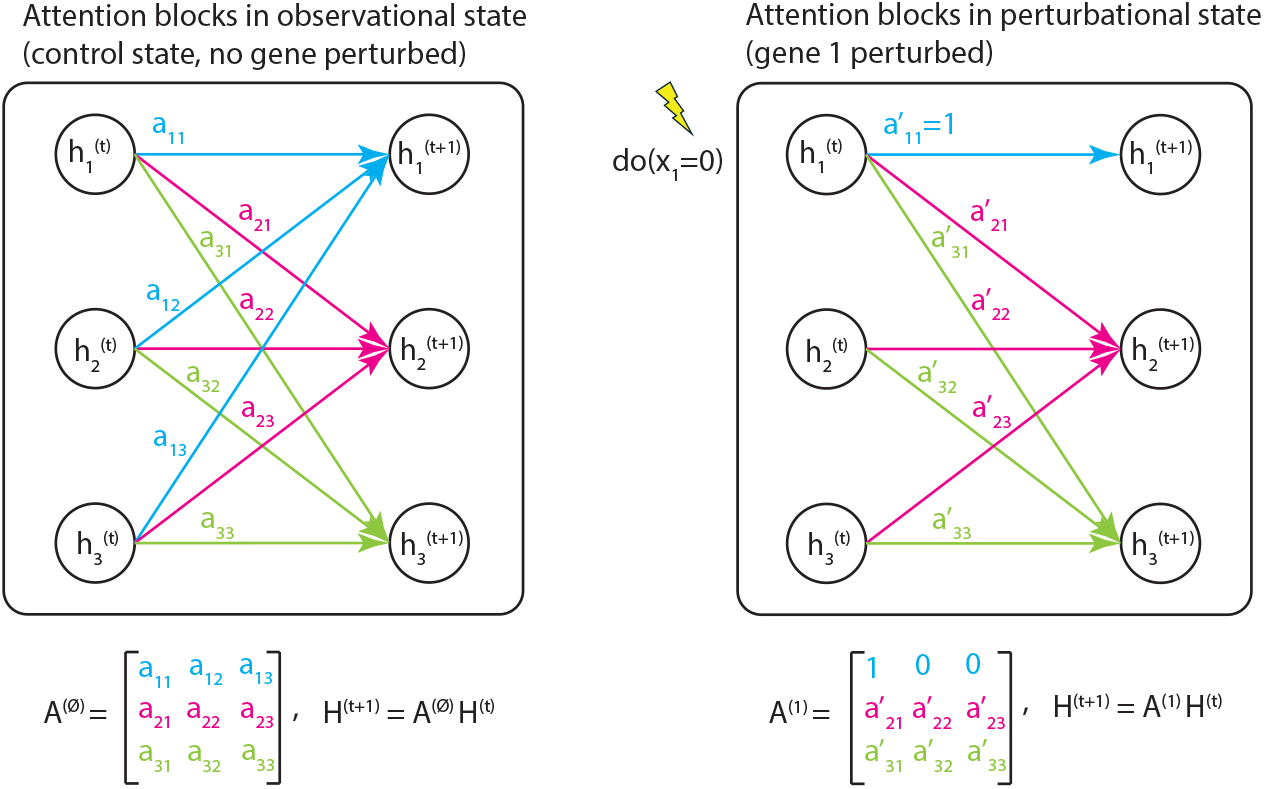
DoFormer’s attention blocks in observational and perturbational states in a simplified three-gene scenario where gene 1 is perturbed.

### 3.3 Input Gene Embeddings

We build the gene embeddings using two complementary sources of information: RNA expression and its protein sequence information. The RNA expression of a gene is cell specific as captured by scRNA-seq data, but its sequence representations are cell agnostic as captured by protein language models (PLM). We use the embeddings of ESM2 (Rives et al. [2021]) or ESMC (Hayes et al. [2025]) model to represent the sequence information of the genes and concatenate it with the RNA expression values to define rich gene embeddings for the model.

Let 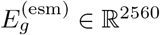 denote the ESM2 embeddings of the gene *g*. In order to make the contribution of RNA expression and ESM2 embeddings equal, we map both to the a *d/*2 dimensional space by learnable linear functions: 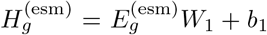, where *W*_1_ ∈ ℝ^2560×*d/*2^ and *b*_1_ ∈ ℝ^*d/*2^, 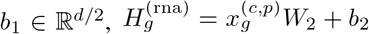, where *W*_2_ ∈ ℝ^1×*d/*2^ and *b*_2_ ∈ ℝ^*d/*2^. The final gene embedding input to the DoFormer model is the concatenation of these two: 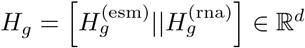.

### 3.4 Training DoFormer

#### 3.4.1 Pre-training on Unperturbed Cells

We first pre-train DoFormer in the observational state on control cells based on MLM loss function. We randomly mask the expression values of 15% of the genes in the input and ask the model to reconstruct them in the output supervised by MSE loss function. We do not expect to learn a causal model in this step, as the nature of data is observational and the model will mainly learn correlative patterns. The main goal of this step is to learn a good initial point for causal learning in the next step. If ℳ = {*g*_1_, …, *g*_*M*_} denotes the set of masked genes in a training batch, the masked input is defined as 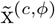, where 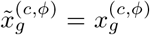, ∀*g* ∈*/* ℳ and 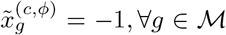, ∀*g* ∈ ℳ. Let 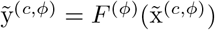 denote the model output. The loss function is defined as MSE on the masked genes: 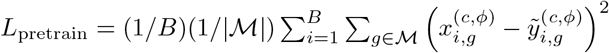, where *B* is the batch size.

#### 3.4.2 Fine-tuning on Perturbed Cells

In the second step, we fine tune the model on the perturbations. This is the core step of enforcing causality in DoFormer model. Before training the model, we first do a differential expression (DE) analysis in the Perturb-seq data to calculate the effect size and DE statistics, which will be used during the training. We first filter the data set to keep only the perturbations with a knock-down efficiency of at least 70% and having at least 30 cells. Then for each perturbation *p* ∈ {1, …, *P*}, we use Wilcoxon rank-sum test to calculate the z-score and p-value for any gene *g* ∈ {1, …, *G*}, denoted respectively by 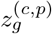 and 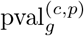. We then define the gene weights 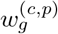 by doing the following steps (similar to Mejia et al. [2025]): 1) absolute value of z-score: 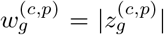, 2) min-max normalization to map them to the range [0, 1]: 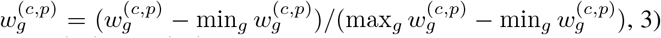 amplify the effects by taking square: 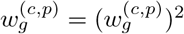, and 4) normalize the weights to sum up to one: 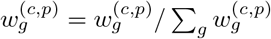. DoFormer will perform two forward passes:

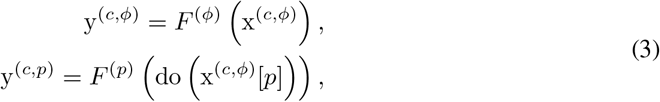

and define Δy^(*c,p*)^ = y^(*c,p*)^ − y^(*c,ϕ*)^. The loss function will be a weighted MSE between predicted delta mean 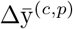 and true delta mean 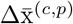 as:

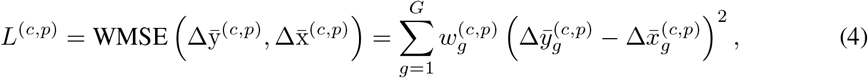

where 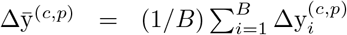 and *B* denotes the batch size. In each training batch, we randomly sample *B* control cells x^(*c,ϕ*)^ from training tuples (*c, p*) ∈ 𝒯 = {(*c*_*i*_, *p*_*j*_)} | cell type c_i_ and perturbation p_j_ in training and compute *L*^(*c,p*)^. The overall loss function is defined as:

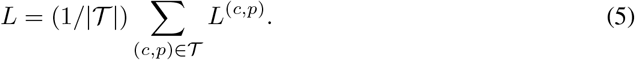

If we have access to only one GPU, in each batch DoFormer updates the gradients based on a single *L*^(*c,p*)^. If we have access to multiple GPU, DoFormer uses Distributed Data Parallel (DDP) where each GPU gets *B* cells for a training tuple (*c, p*), and the gradient updates are based on an average of multiple *L*^(*c,p*)^.

## 4 Results

### 4.1 Train, Validation, and Test Splits

The ultimate goal of a genetic perturbation model is to predict the outcome of gene expression in unseen perturbations in any cell type. However, some of these perturbations may have been observed in other cell types. Let (*c, p*) denote a tuple for perturbation *p* ∈ {1, …, *P*} in the cell type *c* ∈ {1, …, *C*}. Similarly to Adduri et al. [2025], we assume a scenario in which we have access to all perturbations in *C* − 1 cell types and a varying fraction of perturbations in the target (test) cell type, and the goal is to predict the outcome for the remaining perturbations in the target cell type. Let *p*_train_*/p*_validation_*/p*_test_ denote the percentages of perturbations in the target cell type used for training, validation, and test, respectively. We always use 10% for validation, and we try different splits between train and test. For example, 30*/*10*/*60 split means 30% (60%) of perturbations in the target cell type are used for training (test). Figure 2 shows a simple scenario with 4 cell types and 6 perturbations. Note that DoFormer can predict perturbations that have never been seen in other cell types, such as *p*_6_ in Fig. 2, which is one of its unique advantages compared to other models.

**Figure 2:**
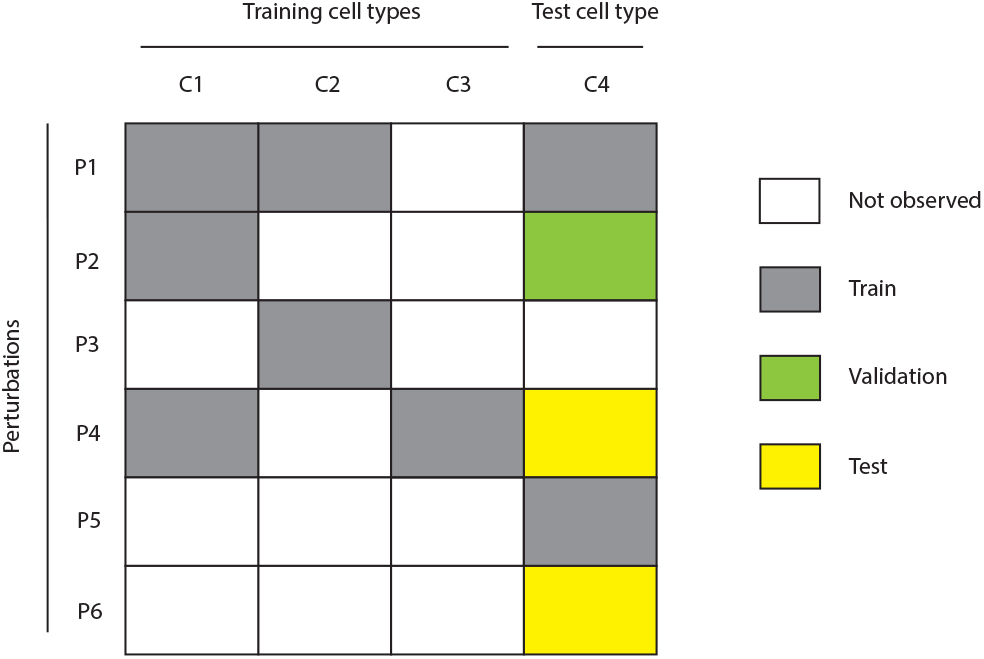
Cell type - perturbation (*c, p*) splits for training, validation, and test sets.

### 4.2 Dataset

Similar to STATE, we combined two large-scale Perturb-seq datasets, Replogle (Replogle et al. [2022]) and Nadig (Nadig et al. [2025]), and built Replogle-Nadig dataset. It consists of four distinct cell types, K562, RPE1, HEPG2, and JURKAT, and 1,855 genetic perturbations after filtering and keeping the perturbations with knockdown efficiency of greater than 70% and with at least 30 cells.

The dataset includes 6,535 genes and 615,810 cells, of which 39,169 are control and 576,641 are perturbed cells. Table 1 shows the numbers per cell type, and the UMAPs are shown in Fig. 6. We were also curious to know in how many cell types each perturbation appears, as shown in Table 2. Note that the perturbations in only one cell type (*N* = 1) are the hardest ones for prediction, as the model has never seen them in other cell types and should do a zero-shot prediction. We expect a causal model to have even bigger performance gap for zero-shot perturbations. The models like STATE (Adduri et al. [2025]) that formulate perturbations based on tokens cannot even predict zero-shot perturbations, which is a big limitation.

**Table 1:** Replogle-Nadig Dataset.

|  | All | K562 | RPE1 | HEPG2 | JURKAT |
| --- | --- | --- | --- | --- | --- |
| # Cells | 615,810 | 180,773 | 168,597 | 91,993 | 174,447 |
| # Cells – control | 39,169 | 10,692 | 11,486 | 4,977 | 12,014 |
| # Cells – perturbed | 576,641 | 170,081 | 157,111 | 87,016 | 162,433 |
| # Genetic Perturbations | 1,855 | 1,312 | 1,451 | 1,251 | 1,437 |

**Table 2:**
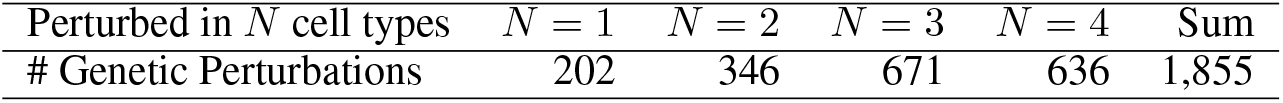
Distribution of Genetic Perturbations Across Cell Types.

### 4.3 Comprehensive Perturbation Prediction Evaluation

We performed a comprehensive evaluation of DoFormer in the Replogle-Nadig dataset across multiple metrics. The complete definitions of these metrics are in the appendix A. In addition to STATE, we also compared DoFormer with two baseline predictors, the perturbation mean (PMean) and the context mean (CMean), as defined in Adduri et al. [2025]. See the appendix C for detailed definitions. As an example, the prediction for (*c*_4_, *p*_4_) in Fig 2 would be as follows: PMean would use the average effect of (*c*_1_, *p*_4_) and (*c*_3_, *p*_4_) (across training cell types), and CMean would use the average effect of (*c*_4_, *p*_1_) and (*c*_4_, *p*_5_) (across training perturbations in test cell type). These simple baselines have been shown to outperform various foundation models in perturbation prediction tasks.

We first pretrained Doformer on control cells from all *C* = 4 cell types. We then fine tuned it four times, each time holding out one target cell type. We started with a 80*/*10*/*10 split in each target cell type. Figure 3 shows that DoFormer significantly outperforms STATE and the baseline models in all metrics in all target cell types. Each point in Fig. 3 is a test perturbation in the target cell type, which is the same for all models. To calculate the significance score, we used Wilcoxon signed-rank test and reported the p-values on top of the distributions (three p-values correspond to DoFormer vs STATE, PMean, and CMean, respectively). Based on these tests, Doformer improvements over STATE, PMean, and CMean are significant in all metrics.

**Figure 3:**
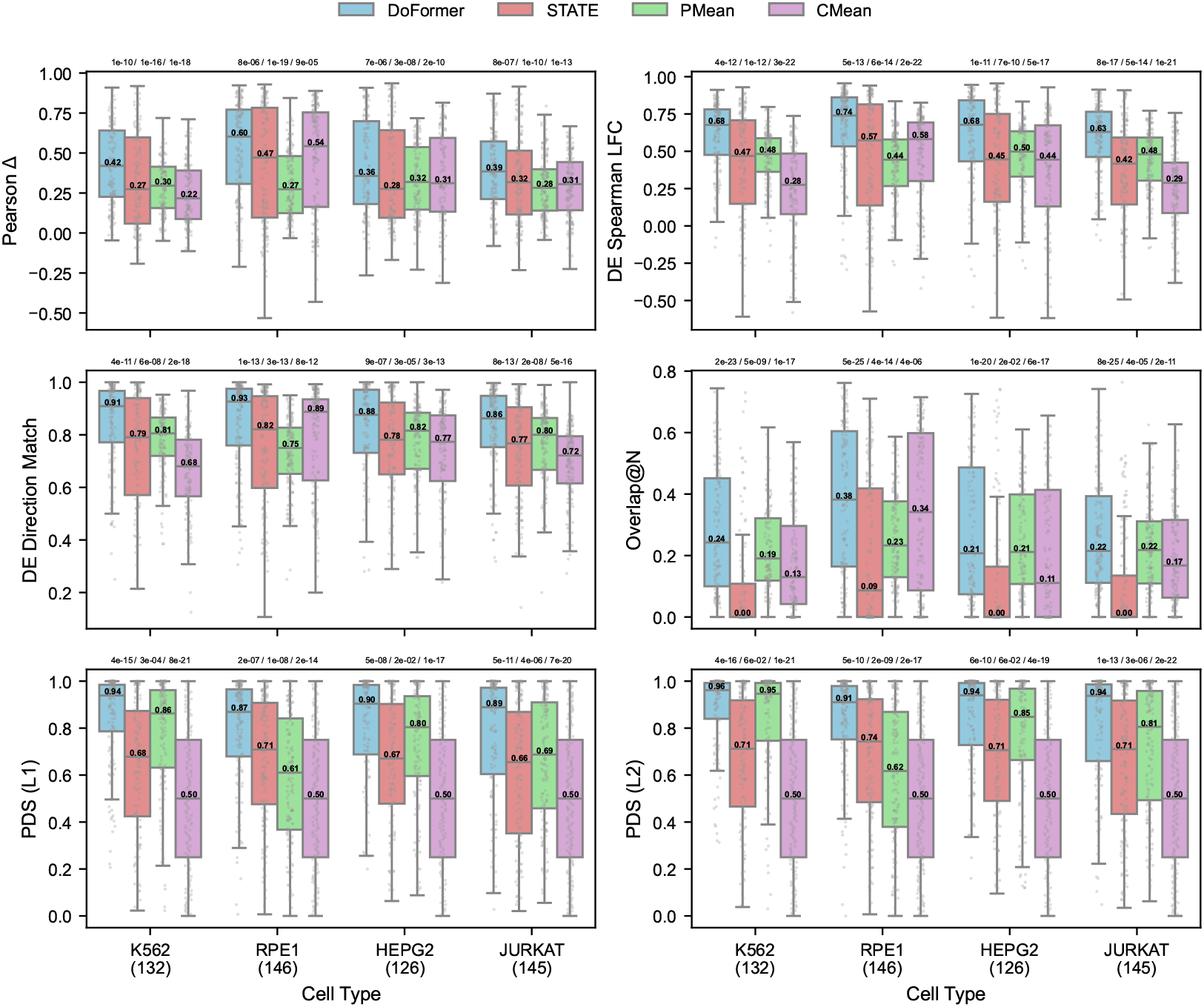
DoFormer significantly outperforms other models across all evaluation metrics and cell types. We use an 80/10/10 split. The number of test perturbations is shown below each target cell type. Each point represents a perturbation, and the median of each distribution is printed. P-values from Wilcoxon signed-rank tests are shown above each distribution, comparing DoFormer vs STATE, PMean, and CMean, respectively. Higher values are better in all metrics.

We find the following observations in Fig. 3 compelling. First, perturbation predictions in some cell types are easier than others; for example, in this case, we see that both DoFormer and STATE have better performance in RPE1 compared to other cell types. This might be because some perturbations create larger changes in gene expression in certain cell types, making them easier for the model to predict. Second, DoFormer outperforms STATE in all-gene metrics (such as Pearson delta) as well as DE-gene metrics; however, the improvement gap is greater for DE-gene metrics such as Overlap_*N*_, for which the median of STATE is zero for three cell types. Third, DoFormer predictions are much more perturbation-specific than STATE, as it gets PDS scores closer to one. DoFormer’s promising performance across all of these metrics comes from its ability to model the *in silico* perturbation correctly. We can also see the pairwise comparisons of DoFormer and STATE for all the perturbations in Figs. 11–14.

Next, we trained the models using different splits: 00*/*10*/*90 (unseen cell type), 10*/*10*/*80, 30*/*10*/*60, 50*/*10*/*40 and 80*/*10*/*10 with K562 as the target cell type. A causal model is expected to behave better by seeing more perturbational data as the size of its equivalence class decreases and can better generalize to unseen perturbations. Since Doformer is a causal model, we expected to see an increasing trend of all the evaluation metrics by expanding the perturbations in training, and that is exactly what we see in Fig. 4. However, STATE does not show an increasing trend in the metrics by expanding train perturbations, especially DE-related metrics like Overlap_*N*_ stay in the median value of zero, suggesting that STATE does not behave as a causal model. DoFormer significantly outperforms the STATE and baseline models in all splits. The only split where DoFormer cannot consistently outperform PMean is 00*/*10*/*90 (unseen cell type). This is in fact an important observation, suggesting that we need at least some perturbations in the target cell types for more accurate perturbation predictions for the remaining genes.

**Figure 4:**
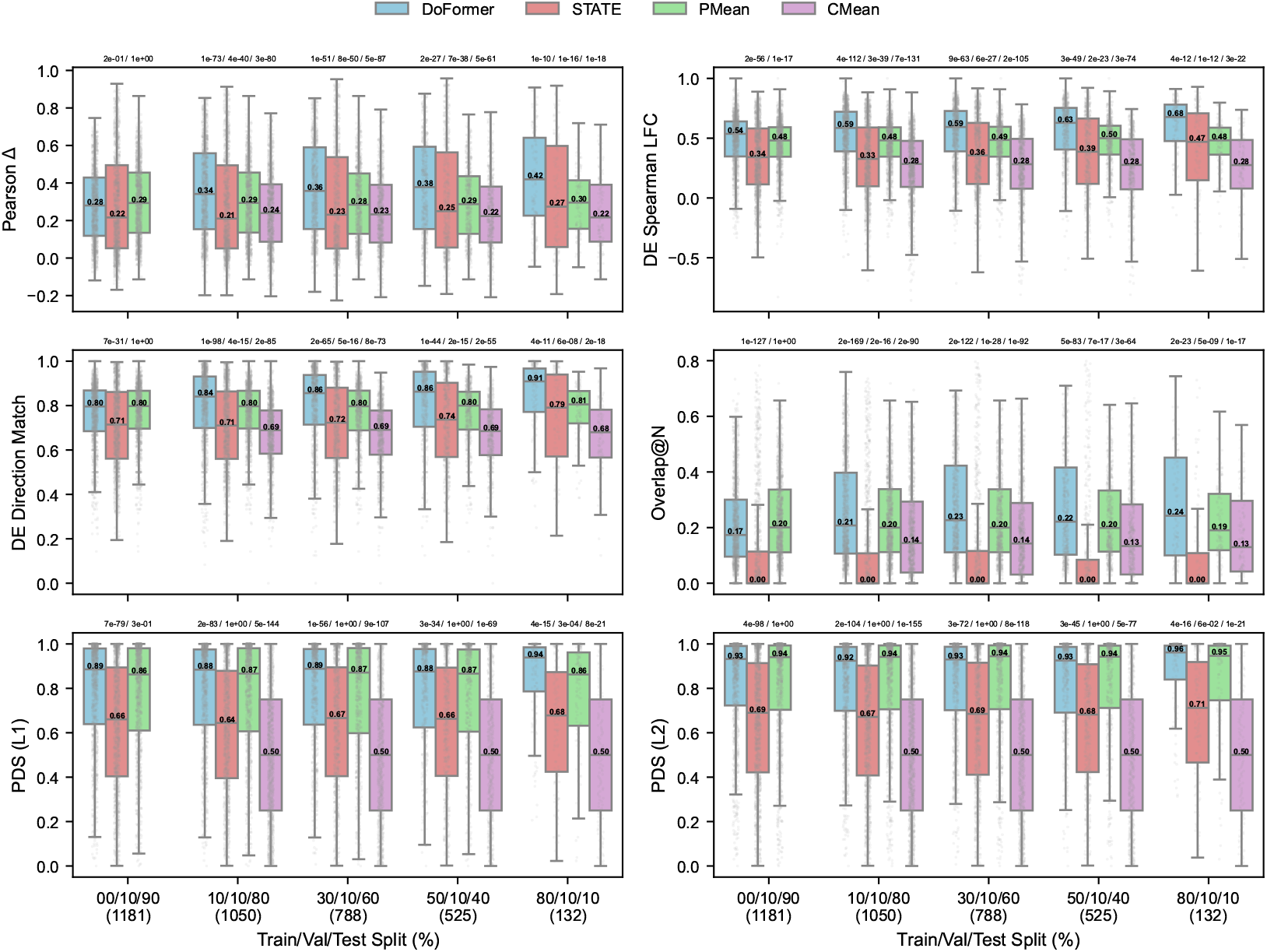
DoFormer significantly outperforms other models across all evaluation metrics and splits. The results are for K562. The number of test perturbations is shown below each split. Each point represents a perturbation, and the median of each distribution is printed. P-values from Wilcoxon signed-rank tests are shown above each distribution, comparing DoFormer vs STATE, PMean, and CMean, respectively. Higher values are better in all metrics.

Finally, we did dive further into the most difficult split 00*/*10*/*90 in K562. This split means that the models have not seen any perturbations in the target cell type K562 during the training. Similar to Table 2, we split the K562 perturbations into four categories of seen in 0 (unseen), 1, 2, and 3 other cell types. Figure 5 shows that DoFormer outperforms STATE in all four categories. We see in Fig. 5 that STATE yields very poor results for the unseen category, where metrics like Overlap_*N*_ is zero for all perturbations, and DE Spearman LFC has a median of 0.14, while DoFormer’s is 0.43. However, DoFormer still yields reasonable metrics in the hardest prediction task of unseen cell type and never seen perturbations. Although DoFormer cannot significantly outperform PMean in all the categories of the split 00*/*10*/*90, it does so for the unseen perturbations, which relate to the hardest task of prediction in unseen cell type and never seen perturbations (like (*c*_4_, *p*_6_) in Fig. 2). DoFormer outperforms STATE and the baseline models in all other splits 10*/*10*/*80 to 80*/*10*/*10 and all categories, shown in the appendix (Figs. 15–18).

**Figure 5:**
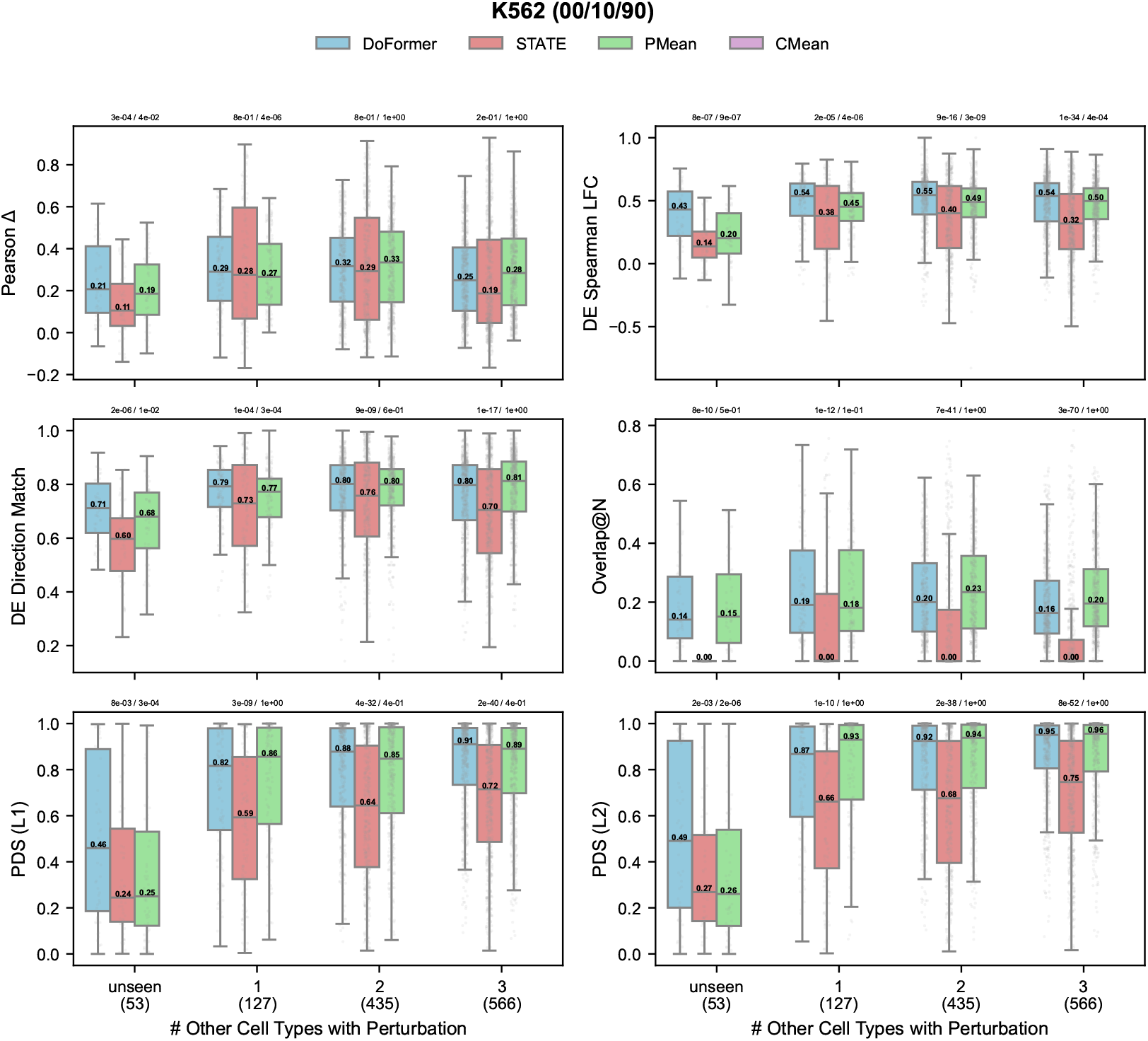
DoFormer outperforms other models in perturbations unseen in other cell types. The results are for K562 in the split 00*/*10*/*90 (zero-shot prediction in K562). The number of test perturbations is shown below each category (unseen, and seen in 1, 2, 3 cell types). Each point represents a perturbation, and the median of each distribution is printed. P-values from Wilcoxon signed-rank tests are shown above each distribution, comparing DoFormer vs STATE and PMean, respectively. Higher values are better in all metrics.

**Figure 6:**
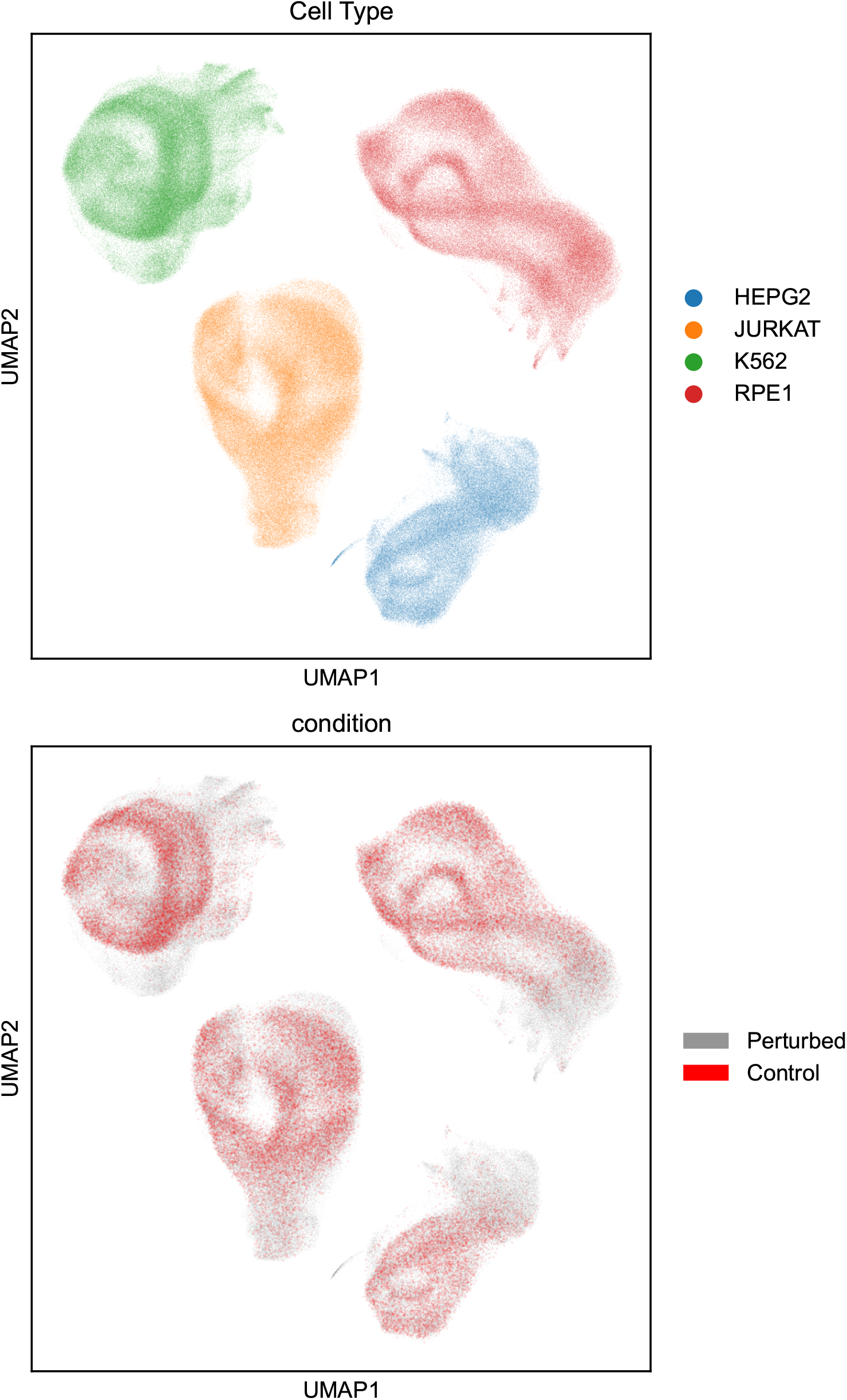
UMAP of Replogle-Nadig dataset with 4 cell types.

Now that we have shown DoFormer to have promising capability in the prediction of genetic perturbations in various metrics, splits, cell types, and categories, we can also investigate its performance visually. Figures 21–28 show the clustermaps of the predicted delta and LFC values for the test perturbations in all four cell types in the split 80*/*10*/*10. We first clustered both rows (perturbations) and columns (genes) using predicted delta (or LFC) and then plotted the heatmap of true delta (or LFC) values using the same indices so that it is easier to compare visually. We can easily see similarity of perturbation-gene block structures between predicted and true delta (or LFC) values, especially for stronger effect sizes, proving that DoFormer can accurately learn the perturbed gene modules for unseen perturbations. See the appendix B for more ablation analyses of DoFormer.

## 5 Conclusions & Limitations

In this work, we introduce DoFormer, a *do*-operator–based Transformer model for *in silico* gene perturbation. DoFormer can be used to generate gene expression outcomes for any genetic perturbation in any cell type by sampling from control cells. We conducted a comprehensive benchmarking using multiple metrics and showed that DoFormer has a promising capability to predict unseen perturbations. DoFormer’s performance mainly stems from *do*-operator–based modifications in the attention blocks. We propose that a causal transcriptomic AI model should distinguish between observational and perturbational data, and DoFormer is a first step in this direction without abandoning scalable Transformer architectures or introducing additional assumptions such as requiring the underlying GRN to be a DAG. One limitation of the current setup is that we assume complete knockdown of perturbed genes (*k* = 0), while this might not always be the case. In future work, we will use the estimated knockdown ratios (0 *< k <* 1) in the CRISPRi data and the activation ratios (*k >* 1) in the CRISPRa data to test whether DoFormer’s generalization ability can be further improved.

## A Comprehensive Evaluation Framework for Perturbation Prediction

What defines a success in a perturbation prediction model? Here we define a set of useful metrics to measure how effective the model predicts the effects of the unseen perturbations. We adopted some of the important CELL-Eval (Adduri et al. [2025]) metrics to our scenario.

Let 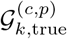 and 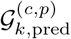 denote the sets of top-*k* DE genes in true and predicted perturbation *p* ∈ {1, …, *P*} in the cell type *c* ∈ {1, …, *C*}, ranked by the absolute delta values 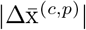 and 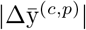, respectively. Let 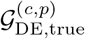 denote the set of all true significant DE genes (*p*_*adj*_ *<* 0.05). To calculate 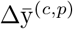 during evaluation, we generate *M* = 1200 perturbed cells for each perturbation in each cell type by randomly sampling *M* control cells from the same cell type.

### A.1 Pearson Delta

This metric measures the Pearson correlation between the true and predicted delta of mean expression values of perturbed and control cells in a given cell type. For a perturbation *p* ∈ {1, …*P*} in a cell type *c* ∈ {1, …, *C*}, it is defined as:

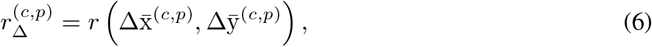

where *r* denotes the Pearson correlation.

### A.2 DE Pearson Delta

This metric is similar to the Pearson Delta but calculated only for the true DE genes for each (*c, p*). It is defined as:

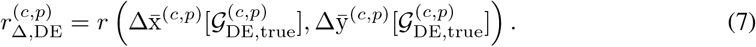

### A.3 DE Spearman Delta

This metric is similar to DE Pearson Delta, but we calculate Spearman’s rank correlation *ρ* instead. It is defined as:

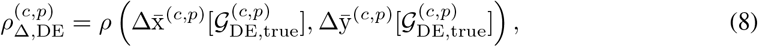

where *ρ* denotes the Spearman correlation.

### A.4 DE Spearman LFC

This metric calculates the Spearman correlation between true and predicted log fold change (LFC) of mean expression values of perturbed and control cells in true DE genes. Since the RNA expression values have undergone the transformation log1p(x) = log(1 + x), we first use the reverse transformation x_count_ = expm1(x) = exp(x) − 1 to bring them back to the count-based space and then calculate the LFC values as

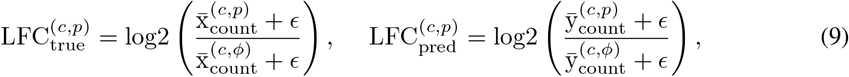

where *ϵ* = 10^−10^, and DE Spearman LFC is defined as

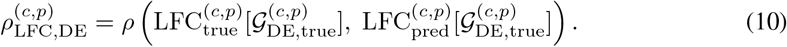

### A.5 DE Directionality Agreement

This metric measures what fraction of true DE genes have the same sign of true and predicted delta values. If this metric is closer to one, it means that the model can correctly predict whether the DE genes will be up- or down regulated after any perturbation. It is defined as:

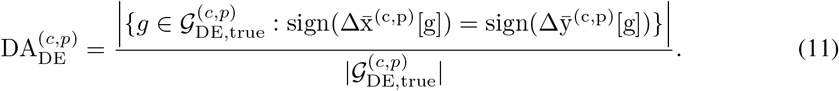

### A.6 Top-*k* Overlap Accuracy

This metric measures overlap between the sets 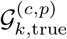 and 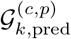 for *k* ∈ {50, 100, 200, *N*}. If *k* = *N*, it means that *k* is equal to the number of true DE genes, that is, 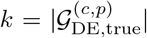, which is perturbation specific. This score is in the range [0, 1], with 1 showing the perfect prediction of top DE genes. It is defined as:

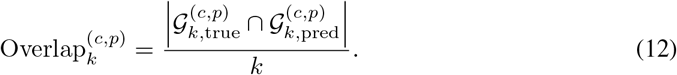

### A.7 AUROC/AUPRC

These metrics measure the area under the receiver operating characteristic (AUROC) and area under the precision-recall curve (AUPRC). The genes in the set 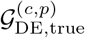 are labeled 1 (significant) and the rest as 0 (non-significant) for any perturbation *p* at the cell type *c*. We use 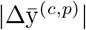 as the confidence score to calculate both metrics. AUROC^(*c,p*)^ evaluates how well the model can distinguish significant from non-significant genes for any perturbation *p* at the cell type *c*. AUPRC^(*c,p*)^ evaluates how well the model can predict the significant genes for any perturbation *p* at the cell type *c* when the number of significant DE genes is low.

### A.8 Perturbation Discrimination Score

This metric measures how perturbation-specific the predictions of the models are. A model works better if it pushes the predicted delta values 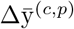 closer to the true delta values 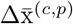 and farther from the other true delta values 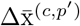, *p*^′^≠ *p*. For any perturbation *p* ∈ {1, …, *P*} in the cell type *c* ∈ {1, …, *C*}, it is defined as:

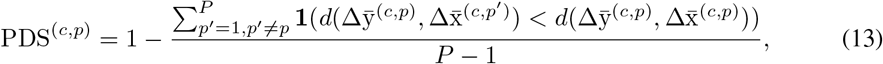

where **1**(.) is the indicator function (1 if true and 0 otherwise), and *d*(.,.) can be one of the following distance functions: *L*_1_ (Manhattan), *L*_2_ (Euclidean), and cosine distance. Note that PDS^(*c,p*)^ ∈ [0, 1], with 1 showing the best discrimination.

## B Ablation Study

Here we ablated different components of DoFormer to understand how they contributed to the predictive performance of the model. We analyzed four main components comparing: 1) PLM embeddings (ESM2 and ESMC) versus non-informative learnable gene embeddings, 2) larger versus smaller model sizes, 3) MSE versus weighted MSE loss functions, and 4) evaluations on all gene set versus highly variable genes (HVG).

### B.1 PLM Embeddings

We first trained two types of small DoFormer models (see Table 3), first with ESM2 (MIT license) embeddings and the second with learnable gene embeddings. We hypothesized that using ESM2 embeddings should provide rich orthogonal information for the genes, helping to learn more causal gene-gene interactions and enhance the predictive power of the model. Figure 7 confirms this hypothesis, as we see that DoFormer with ESM2 embeddings significantly outperforms DoFormer without those embeddings in all of the evaluation metrics. The results of Fig. 7 are for K562 in the split 80*/*10*/*10. We then trained three types of larger DoFormer models (see Table 3), first with ESM2 embeddings, the second with ESM-C (Forge API license) embeddings, and the third with learnable gene embeddings (no ESM). ESMC is the newest model of the ESM family, which outperforms ESM2 in tasks related to protein structures and functions. Interestingly, in the case of large DoFormer models, using ESM embeddings (either ESM2 or ESMC) does not significantly improve performance in all metrics, as shown in Fig. 8. One possible explanation for this behavior could be that adding additional information from the PLM embeddings to the smaller models helps them learn causal interactions; however, the larger models seem to have the capacity to learn causal interactions without any extra information from the PLM embeddings. Since we have no prior idea about which model sizes prevent the need for PLM embeddings in each dataset, we recommend using them as a default in DoFormer.

**Table 3:** Model size configurations in DoFormer.

|  | DoFormer-Small | DoFormer-Large |
| --- | --- | --- |
| # Transformer layers ($L$) | 4 | 8 |
| # Attention heads ($H$) | 4 | 8 |
| Model dimension ($d_{model}$) | 128 | 512 |
| Feed-forward dimension ($d_{ff}$) | 1024 | 2048 |
| # Parameters (approx.) | 1.7M | 27M |

**Figure 7:**
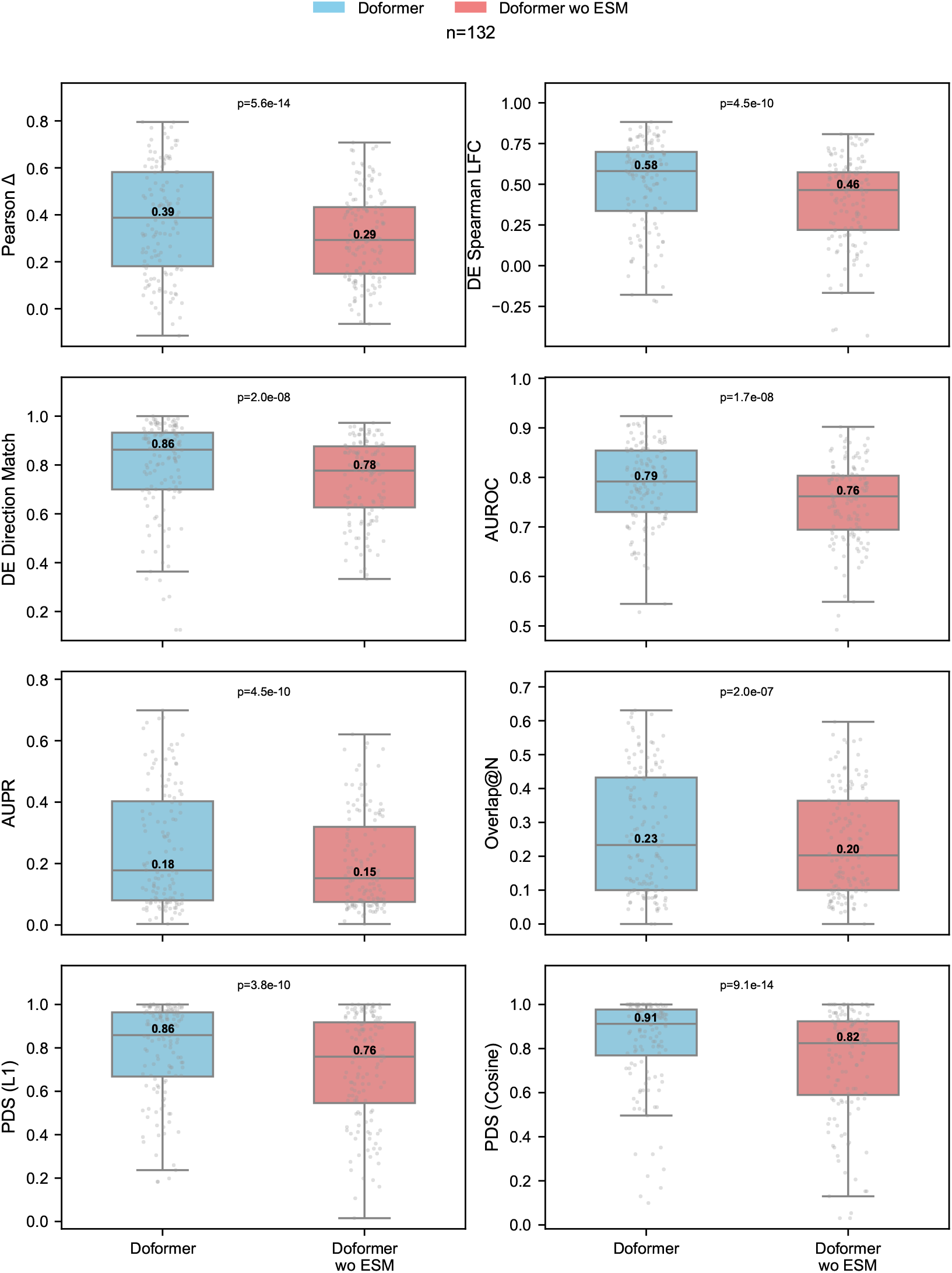
Small DoFormer model with ESM2 embeddings significantly outperforms small DoFormer model without ESM2 embeddings across all evaluation metrics. The results are for K562, small model, and split 80*/*10*/*10. Each point represents a perturbation, and the median of each distribution is printed. P-values from Wilcoxon signed-rank tests are shown above each distribution.

**Figure 8:**
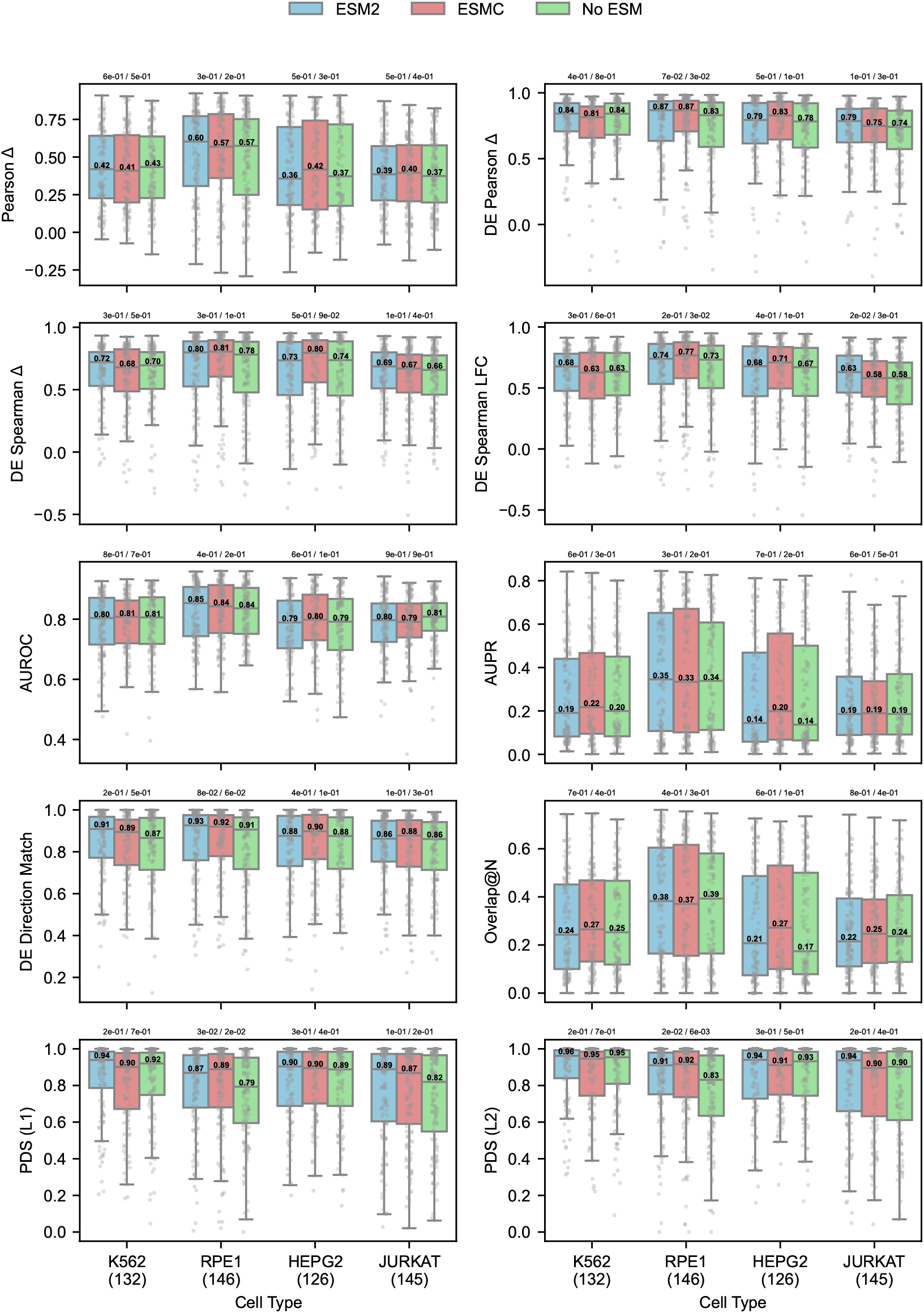
Large DoFormer model with ESM2/ESMC embeddings do not significantly outperform large DoFormer model without ESM embeddings across all evaluation metrics. The results are for K562 and split 80*/*10*/*10. Each point represents a perturbation, and the median of each distribution is printed. P-values from Wilcoxon signed-rank tests are shown above each distribution, comparing ESM2 vs ESMC and No-ESM, respectively.

### B.2 Model Size

We investigated the role of model size on its performance. To this end, we trained two DoFormer models: small and large. The difference is in the number of Transformer layers (*L*), heads (*H*), model dimension *d*_*model*_ and feed-forward dimension (*d*_*ff*_). The small model has *L* = *H* = 4, *d*_*model*_ = 128, and *d*_*ff*_ = 1024, with approximately 1.7*M* parameters. The large model has *L* = *H* = 8, *d*_*model*_ = 512, and *d*_*ff*_ = 2048, with approximately 27*M* parameters (Table 3). Figure 9 shows that the large model significantly outperforms the small model in all evaluation metrics in all cell types. This is an interesting observation, as it is not trivial to improve the model performance across all metrics consistently by only scaling. However, it is possible in DoFormer due to its inherent capability in learning causal gene-gene interactions, which makes it benefit from model scaling. All the results in the paper are from the larger DoFormer model by default.

**Figure 9:**
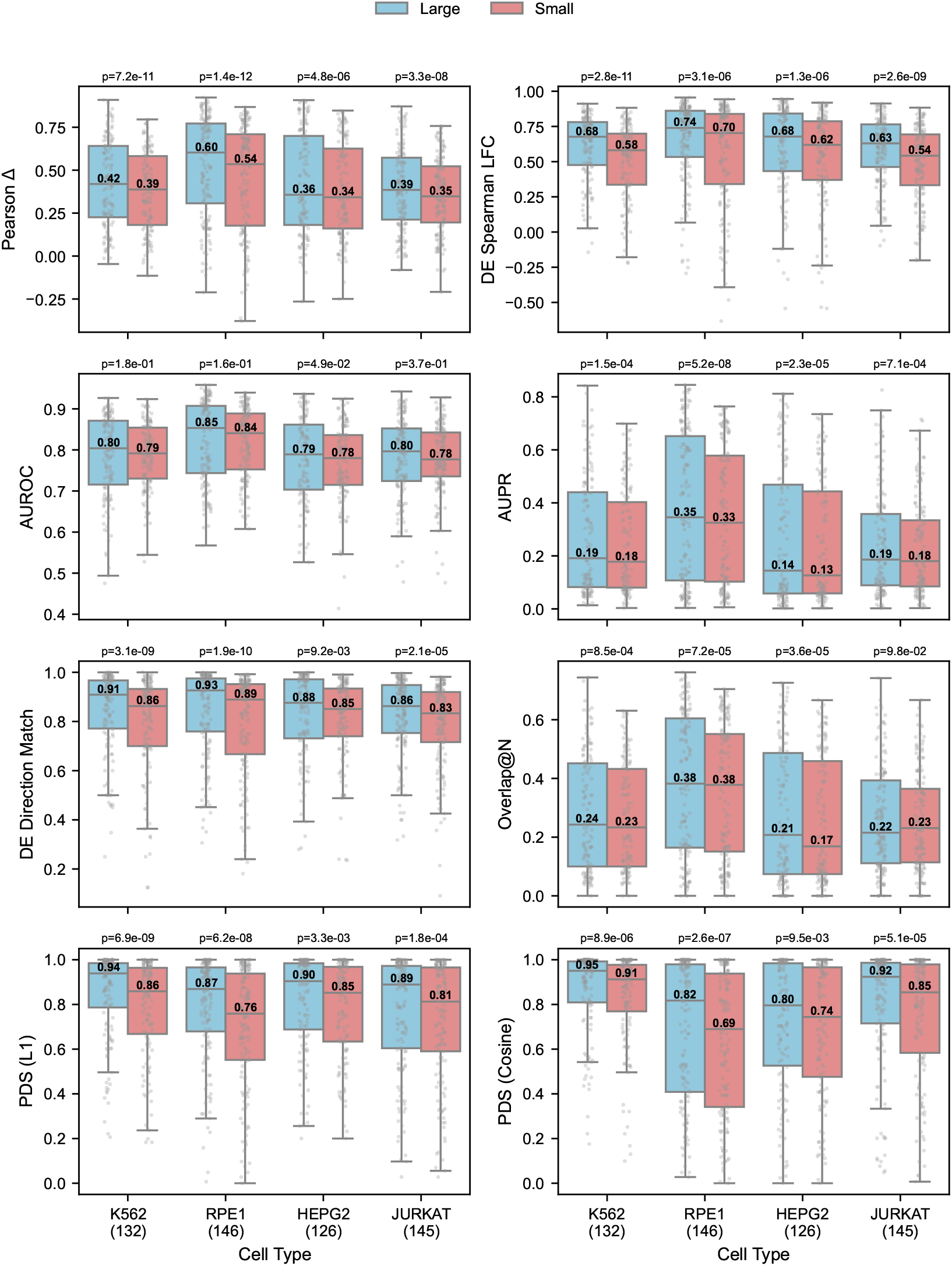
Large DoFormer model significantly outperforms small DoFormer model across all evaluation metrics and cell types. The results are for split 80*/*10*/*10. The number of test perturbations is shown below each cell type. Each point represents a perturbation, and the median of each distribution is printed. P-values from Wilcoxon signed-rank tests are shown above each distribution.

### B.3 Loss Function

Although we used a weighted MSE loss function in DoFormer to encourage prioritization of DE genes, we were interested in understanding how the model would work compared to MSE loss function without using weights. Figure 10 shows this comparison in all cell types for the split 80*/*10*/*10. It is interesting to see that unlike the model size, there is no clear winner in this case, as it is metric dependent. MSE works better in evaluation metrics that consider all genes, such as Pearson Delta, and WMSE significantly outperforms MSE in DE-related metrics, such as DE Spearman LFC. It is also interesting that using WMSE leads to significantly higher perturbation discrimination scores, which suggests WMSE help the models escape the mode collapse and generate perturbation-specific predictions. Based on these results, we recommend WMSE over MSE, unless one really cares about Pearson Delta, in which case MSE sounds a better choice.

**Figure 10:**
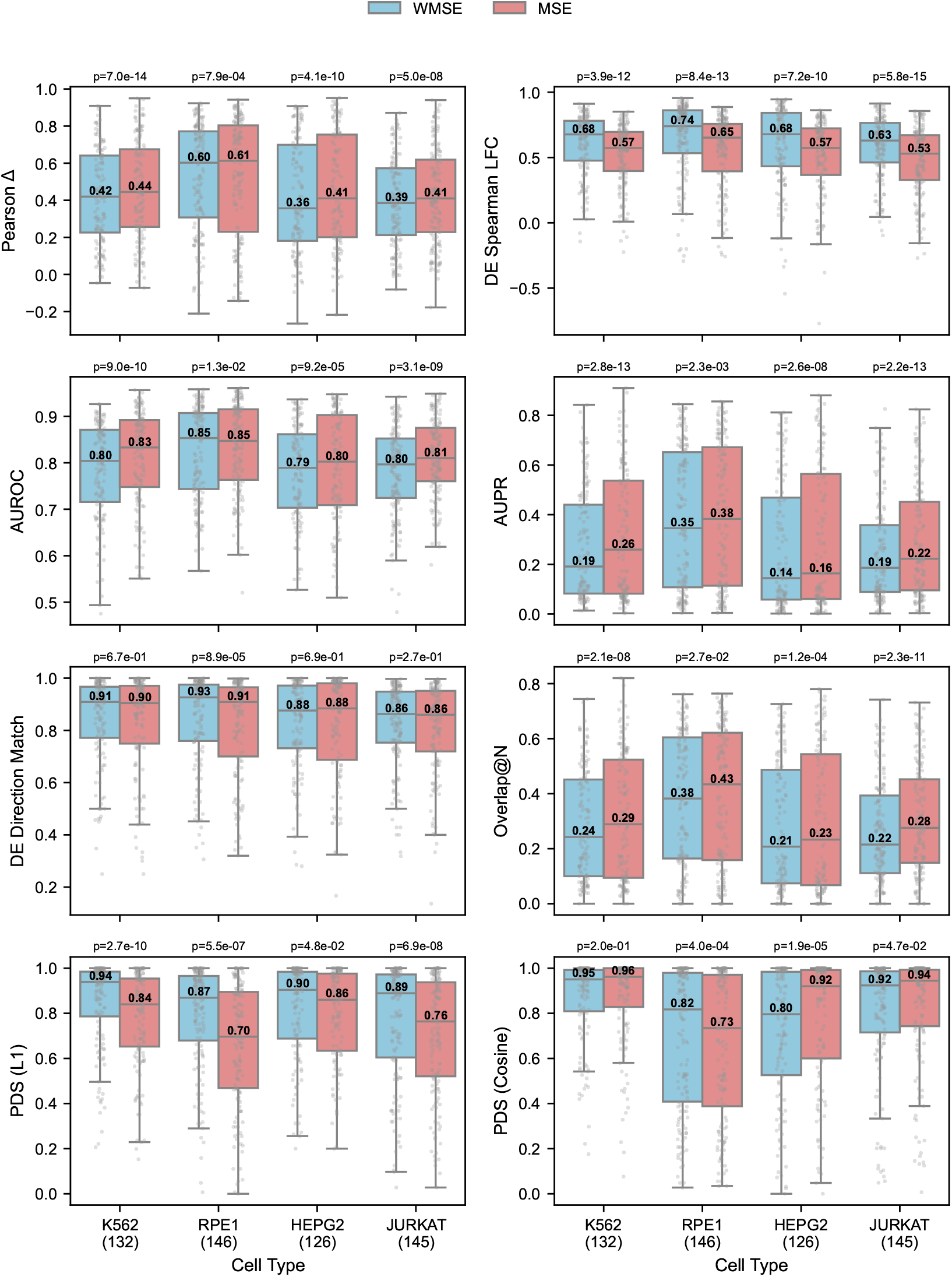
WMSE versus MSE loss function in DoFormer’s performance. WMSE leads to better DE-related metrics, while MSE leads to better all-gene metrics like Pearson Delta. Neither loss function outperforms the other in all metrics. The results are for split 80*/*10*/*10. The number of test perturbations is shown below each cell type. Each point represents a perturbation, and the median of each distribution is printed. P-values from Wilcoxon signed-rank tests are shown above each distribution.

### B.4 All versus Highly Variable Genes

Usually foundation models use 2000 highly variable genes (HVGs) as the feature set to reduce the complexity of the model. However, it is useful to predict the effects of genetic perturbations on the set of all genes. That is why we used all genes in the Replogle-Nadig dataset (6535 genes after merging and filtering) in DoFormer models and performed all evaluation benchmarks based on all genes. Here, we subset the genes to HVGs and performed the same evaluations on this subset to see how they compare to the full gene set. Figure 19 compares the evaluation metrics of DoFormer and baseline models (PMean and CMean) using all genes versus HVGs in the split 80*/*10*/*10 in all target cell types. Figure 20 also compares them in all the splits (00*/*10*/*90 to 80*/*10*/*10) of K562 cells. In general, we can see that the evaluation metrics are higher (better) using HVGs (compared to all genes), especially in DE-related metrics. This makes intuitive sense since HVGs are a smaller set of genes, and accurate prediction of perturbation outcomes in all genes is more difficult.

## C Baseline Models

We consider two baseline models for perturbation predictions that have been shown to sometimes outperform foundation models: the perturbation mean (PMean) and the context mean (CMean).

### C.1 Perturbation Mean (PMean) Baseline

Assume that we want to predict the effect of perturbation *p* ∈ {1, …, *P*} in the test cell type *c*_*t*_ ∈ {1, …, *C*}. PMean uses the average of delta values for the perturbation *p* seen in the cell types other than *c*_*t*_. Let C_*p*_ denote the set of all the cell types in training data that have the perturbation *p*. If |C_*p*_| *>* 0, PMean predictor for the test cell type *c*_*t*_ and the perturbation *p* is defined as:

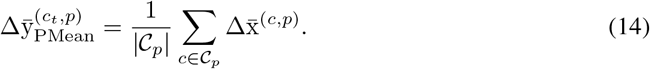

If |𝒞_*p*_| = 0 (perturbation in the test cell type never seen in other cell types), we define PMean as the average of all training (*c, p*) pairs in all cell types except *c*_*t*_:

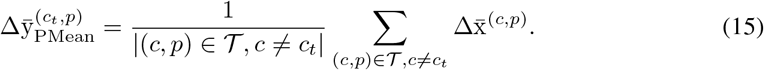

### C.2 Context Mean (CMean) Baseline

Let assume we want to predict the effect of perturbation *p* ∈ {1, …, *P*} in the target cell type *c*_*t*_ ∈ {1, …, *C*}. CMean uses the delta of control cells and all perturbed cells in the training data in the target cell type:

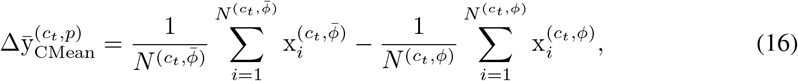

where 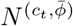 denotes the number of all perturbed cells of the cell type *c*_*t*_ in the training set. Note that CMean uses the same delta prediction for all test perturbations, and CMean is not defined for the split 00*/*10*/*90 since there are no perturbed cells in the training set for the target cell type 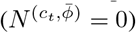.

## D Compute Resources

We have trained DoFormer models in an SLURM-based HPC cluster with access to H100 (80G VRAM) and H200 (140G VRAM) GPUs, where each compute node has 8 GPUs. We have always trained the models on one node (either H100 or H200) and usually distributed over 4 GPUs using DDP. For each run, we also requested 16 CPUs per task and 200G RAM per GPU (800G overall). As such, in the Replogle-Nadig dataset, pre-training took almost 15 hours and fine-tuning almost 1 day.

## E Training Details and Hyperparameters

We used the batch size of 32 cells per GPU during pre-training DoFormer. With 4 GPUs, the effective batch size would be 128 cells. We used 90% of the control cells for training and 10% for validation to monitor the MSE loss functions. For fine-tuning, we used the batch size of 24 cells per GPU, with the effective batch size of 96 cells with 4 GPUs. The train, test, and validation splits have been explained in the paper. We saved the model checkpoints every 500 steps and recorded the validation loss every 500 steps. We used an early stopping criterion with patience of 20, meaning that the model will stop training if there is no improvement over 10,000 steps. We used AdamW optimizer with the base learning rate of 1*e* − 4 and weight decay of 1*e* − 5. We adopted a cosine annealing with a warm-up strategy for the learning rate, where it increases linearly from zero to the maximum value of 1*e* − 4 in 1000 steps and decreases to the minimum value of 1*e* − 5 in 100 epochs (maximum epoch size) following a cosine function.

Regarding the STATE model, we used their ST model and followed their code at https://github.com/ArcInstitute/state (CC BY-NC-SA 4.0 license) to train them from scratch on exactly the same training, validation, and test sets that we used in DoFormer to make comparisons fair.

## F Dataset

The Replogle-Nadig dataset consists of 4 cell types. K562 and RPE1 are from the Replogle dataset (Replogle et al. [2022]), which can be downloaded from this link: Figshare (CC BY 4.0 license). HEPG2 and JURKAT are from the Nadig dataset (Nadig et al. [2025]), which can be downloaded from this link: GEO. We merged these two datasets and performed the basic preprocessing of the single cells. We normalized the cells to have total counts of 10,000 and applied the *log*1*p* transformations to all expression values. For filtering perturbations, we followed the steps of cell-load package (cell-load) with the default values of residual_expression = 0.30 (meaning 70% knockdown efficiency), cell_residual_expression = 0.50 (cell-level threshold), and min_cells = 30 (minimum cells per perturbation after filtering). We only kept the perturbations that are within the full gene set after merging the datasets since we need them to be in the DoFormer model for the *in silico* perturbations.

## G Code Availability

The DoFormer code will be made available to the public after the publication of the manuscript.

## H Broader Impacts

This paper introduces a novel model, DoFormer, for the prediction of genetic perturbations. DoFormer can have multiple positive societal impacts in curing or preventing diseases. Biologists and scientists might find it very helpful in finding the target genes that cause various diseases without running millions of costly experiments in the labs. DoFormer can be used to perform low-cost *in silico* perturbations across all human cells and diseases to discover the causal genes driving each disease. Then, the top predicted targets can be further validated in the experimental labs with a fraction of the cost. Regarding the negative societal impacts, like any biological AI model, DoFormer can also carry inherent dual-use risks such as the theoretical potential to identify vulnerabilities for harmful cellular manipulation.

---

**Figure 11:**
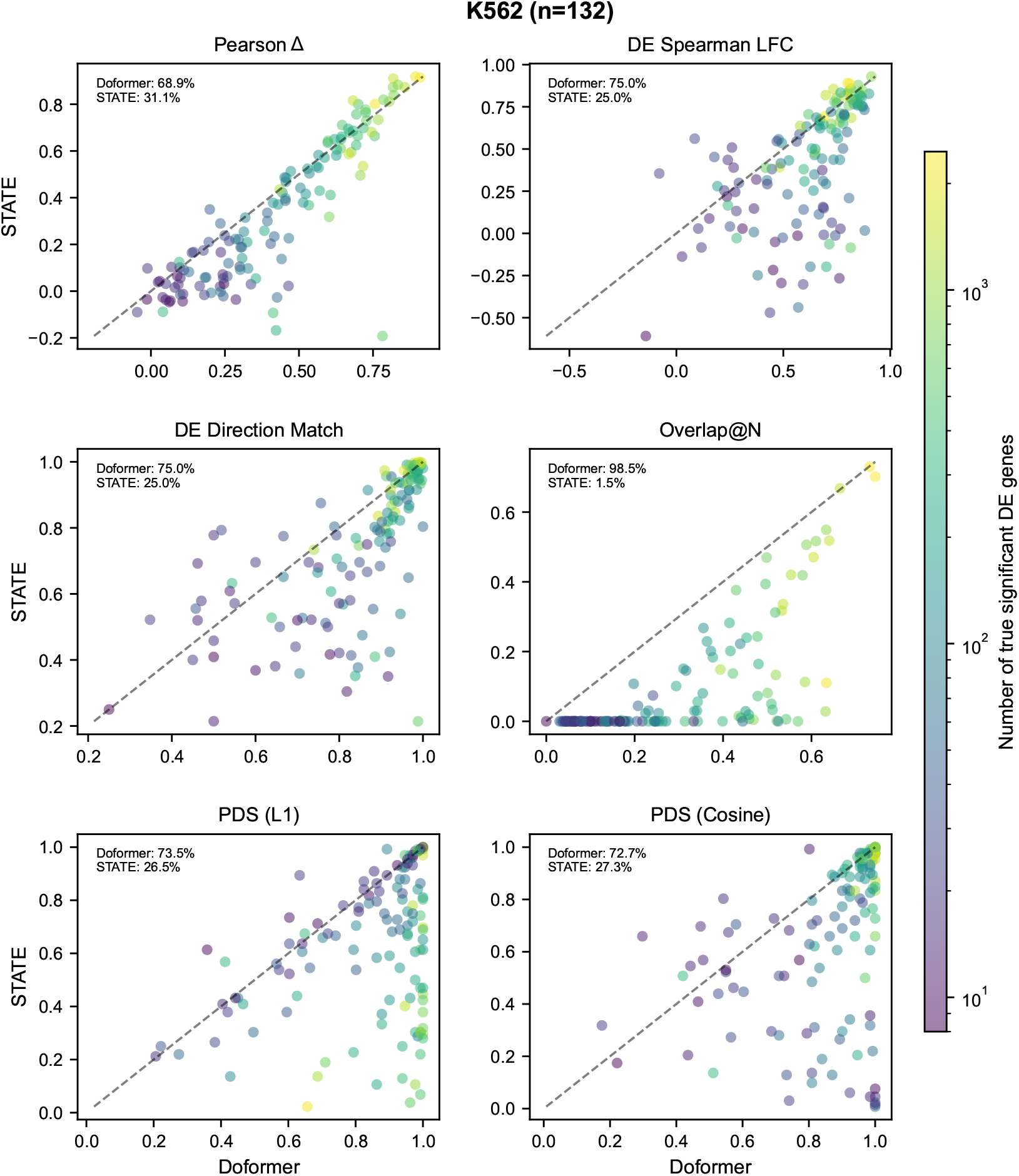
Scatter plots of evaluation metrics comparing DoFormer versus STATE for each test perturbation. The results are for K562 and split 80*/*10*/*10. For the perturbations below the diagonal line, DoFormer’s performance is better than STATE. The percentages of the perturbations for which either model works better are shown in each plot. The perturbations are colored by their true significant DE genes (effect size).

**Figure 12:**
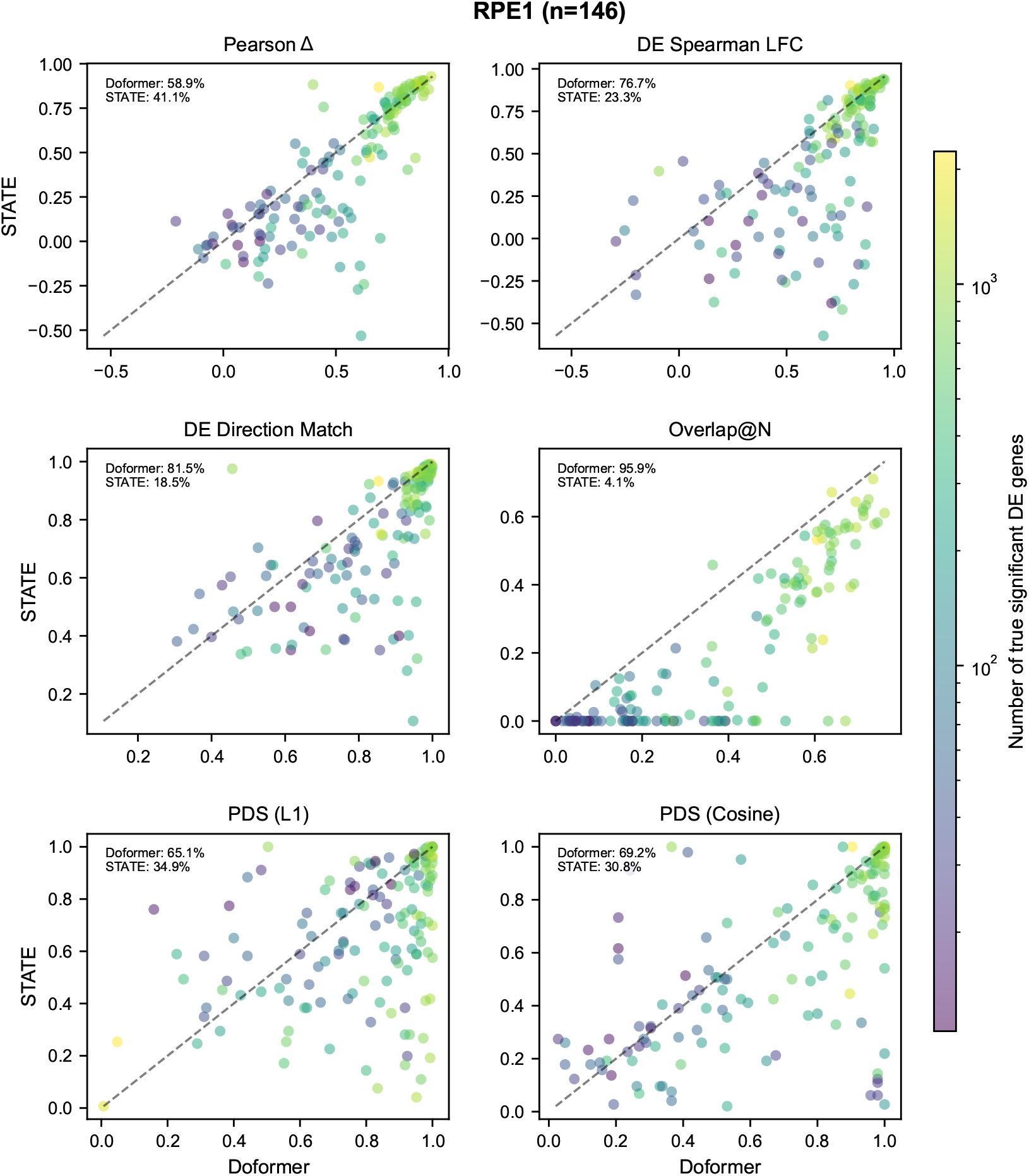
Scatter plots of evaluation metrics comparing DoFormer versus STATE for each test perturbation. The results are for RPE1 and split 80*/*10*/*10. For the perturbations below the diagonal line, DoFormer’s performance is better than STATE. The percentages of the perturbations for which either model works better are shown in each plot. The perturbations are colored by their true significant DE genes (effect size).

**Figure 13:**
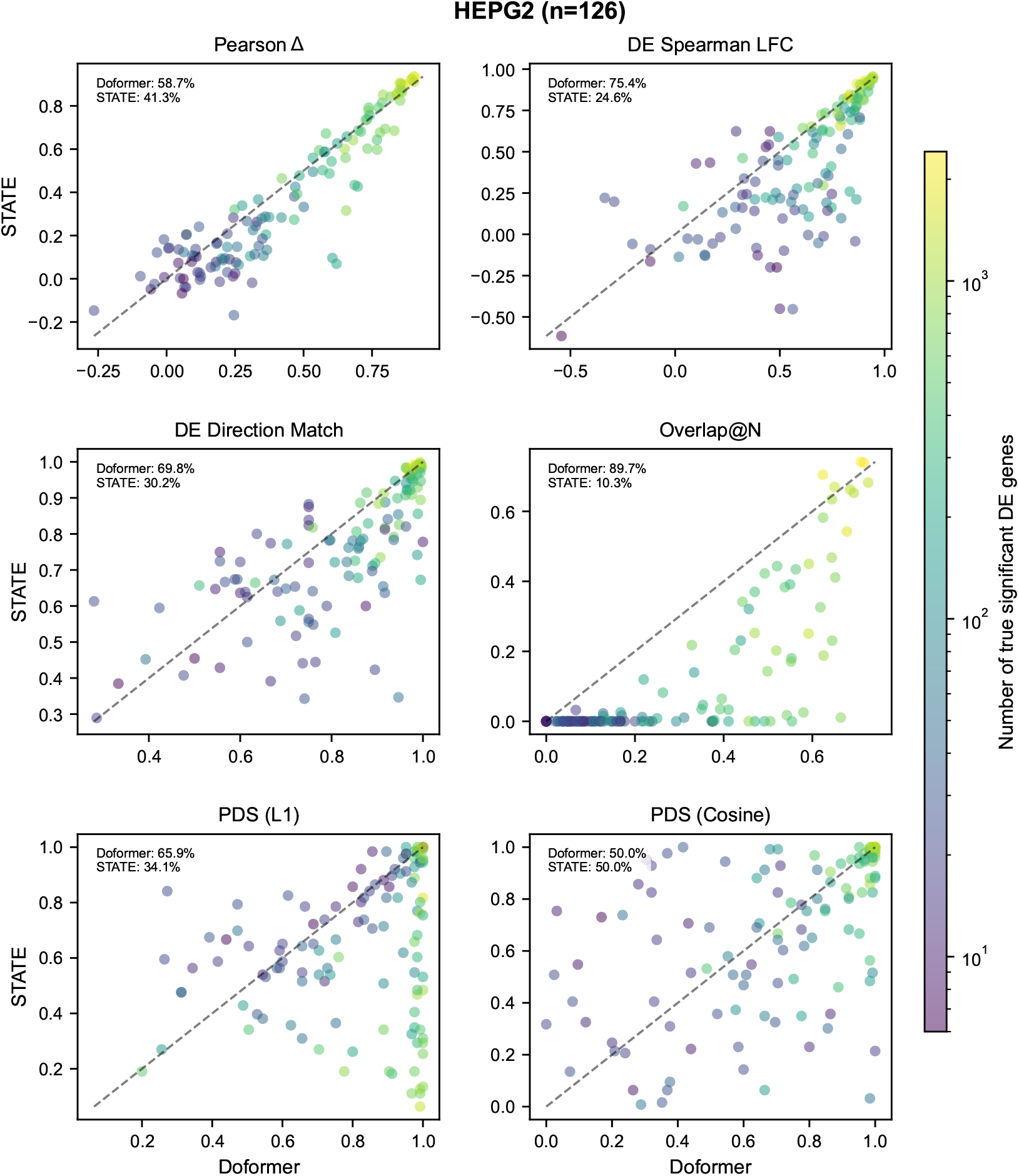
Scatter plots of evaluation metrics comparing DoFormer versus STATE for each test perturbation. The results are for HEPG2 and split 80*/*10*/*10. For the perturbations below the diagonal line, DoFormer’s performance is better than STATE. The percentages of the perturbations for which either model works better are shown in each plot. The perturbations are colored by their true significant DE genes (effect size).

**Figure 14:**
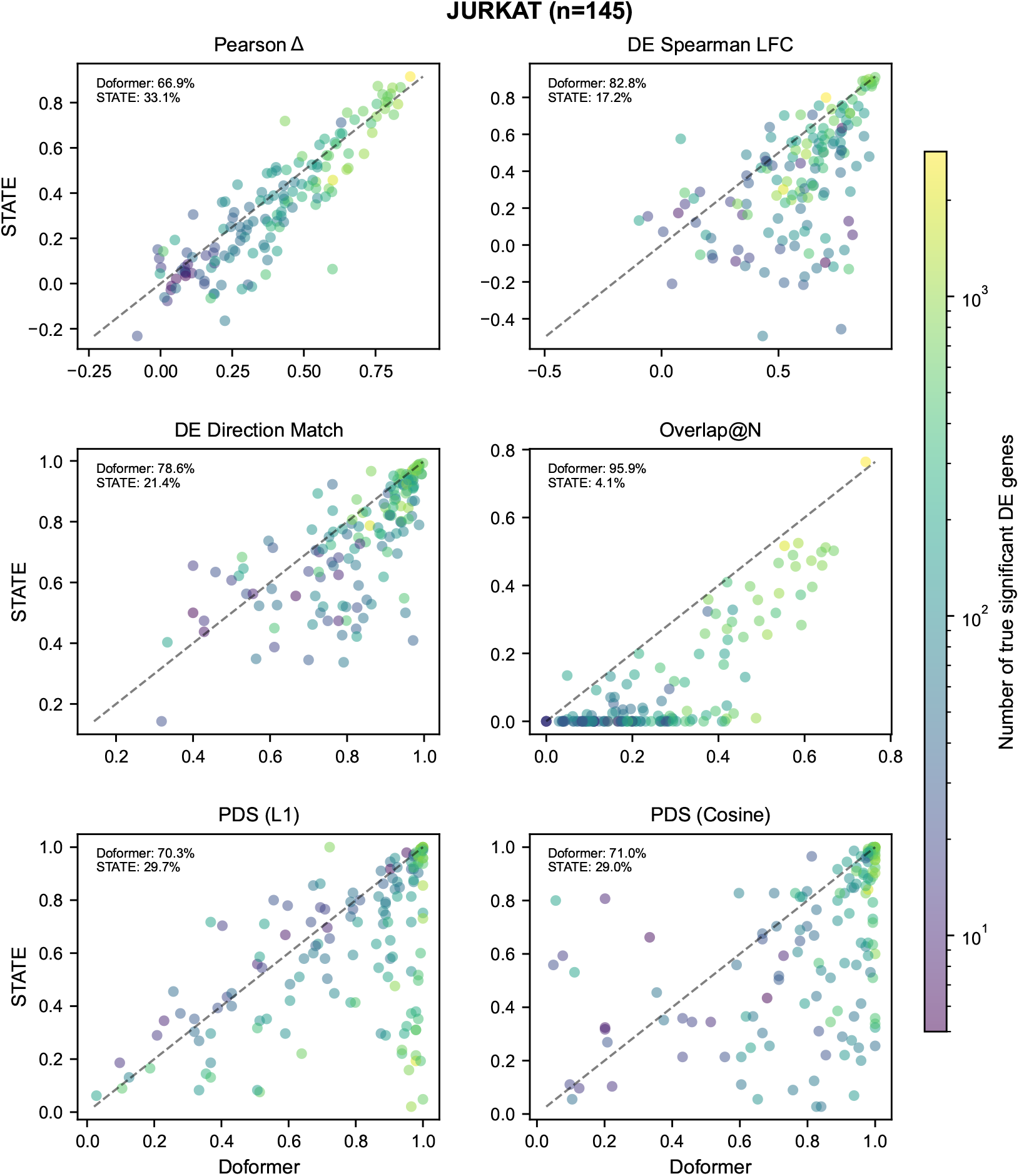
Scatter plots of evaluation metrics comparing DoFormer versus STATE for each test perturbation. The results are for JURKAT and split 80*/*10*/*10. For the perturbations below the diagonal line, DoFormer’s performance is better than STATE. The percentages of the perturbations for which either model works better are shown in each plot. The perturbations are colored by their true significant DE genes (effect size).

**Figure 15:**
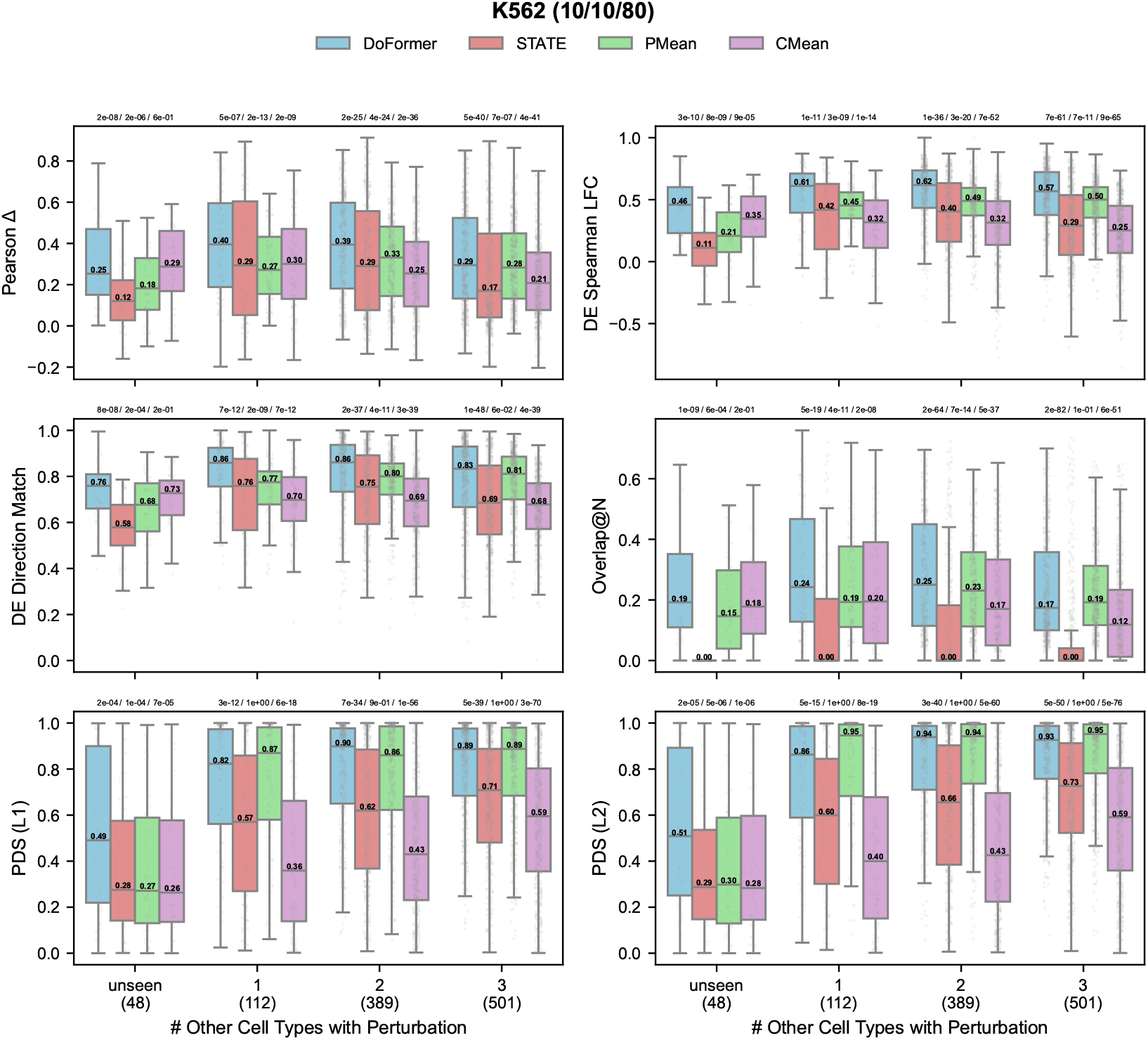
DoFormer outperforms STATE regardless of perturbations seen or unseen in other cell types. The results are for K562 in the split 10*/*10*/*80. The number of test perturbations is shown below each category (unseen, and seen in 1, 2, 3 cell types). Each point represents a perturbation, and the median of each distribution is printed. P-values from Wilcoxon signed-rank tests are shown above each distribution.

**Figure 16:**
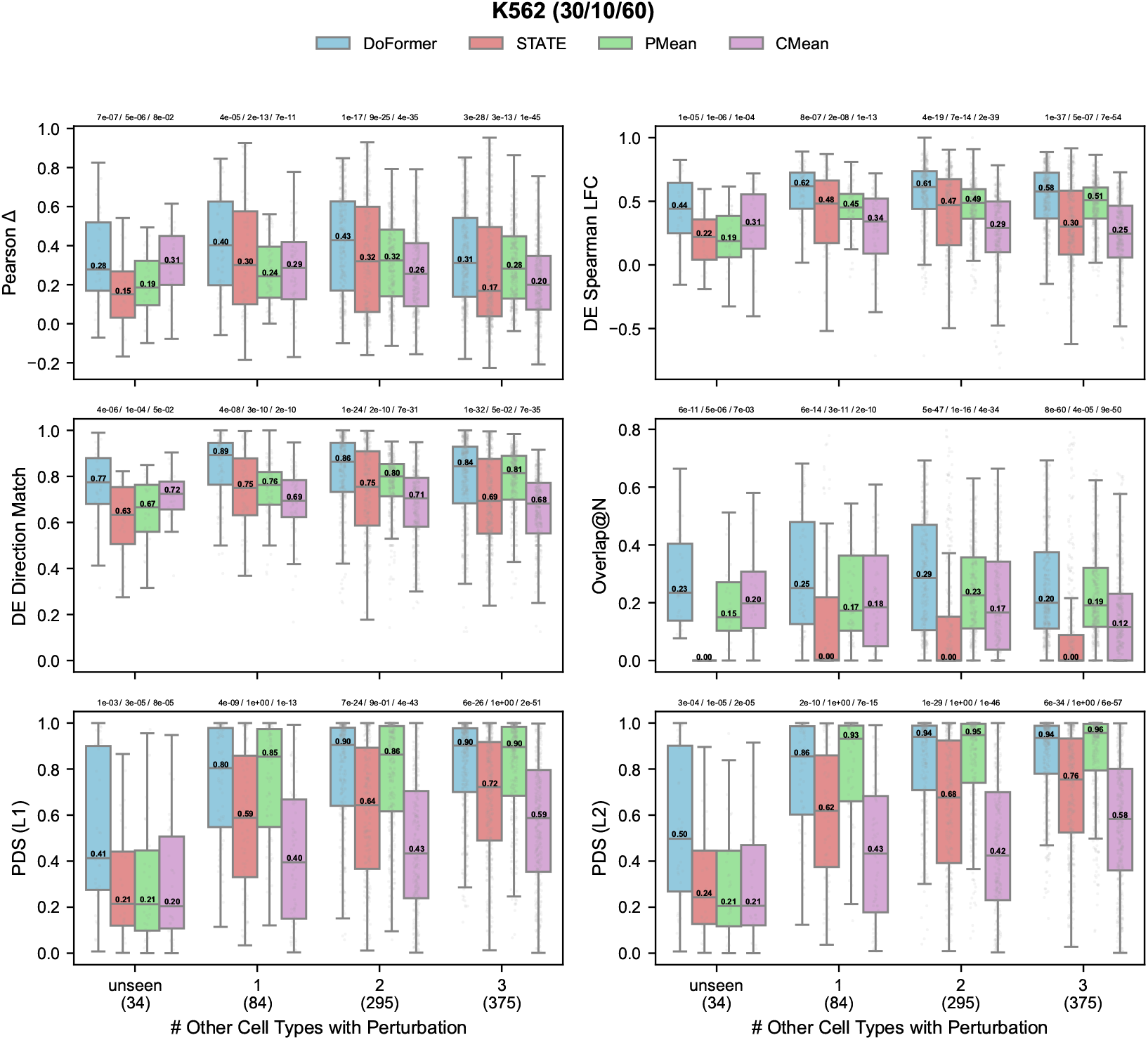
DoFormer outperforms STATE regardless of perturbations seen or unseen in other cell types. The results are for K562 in the split 30*/*10*/*60. The number of test perturbations is shown below each category (unseen, and seen in 1, 2, 3 cell types). Each point represents a perturbation, and the median of each distribution is printed. P-values from Wilcoxon signed-rank tests are shown above each distribution.

**Figure 17:**
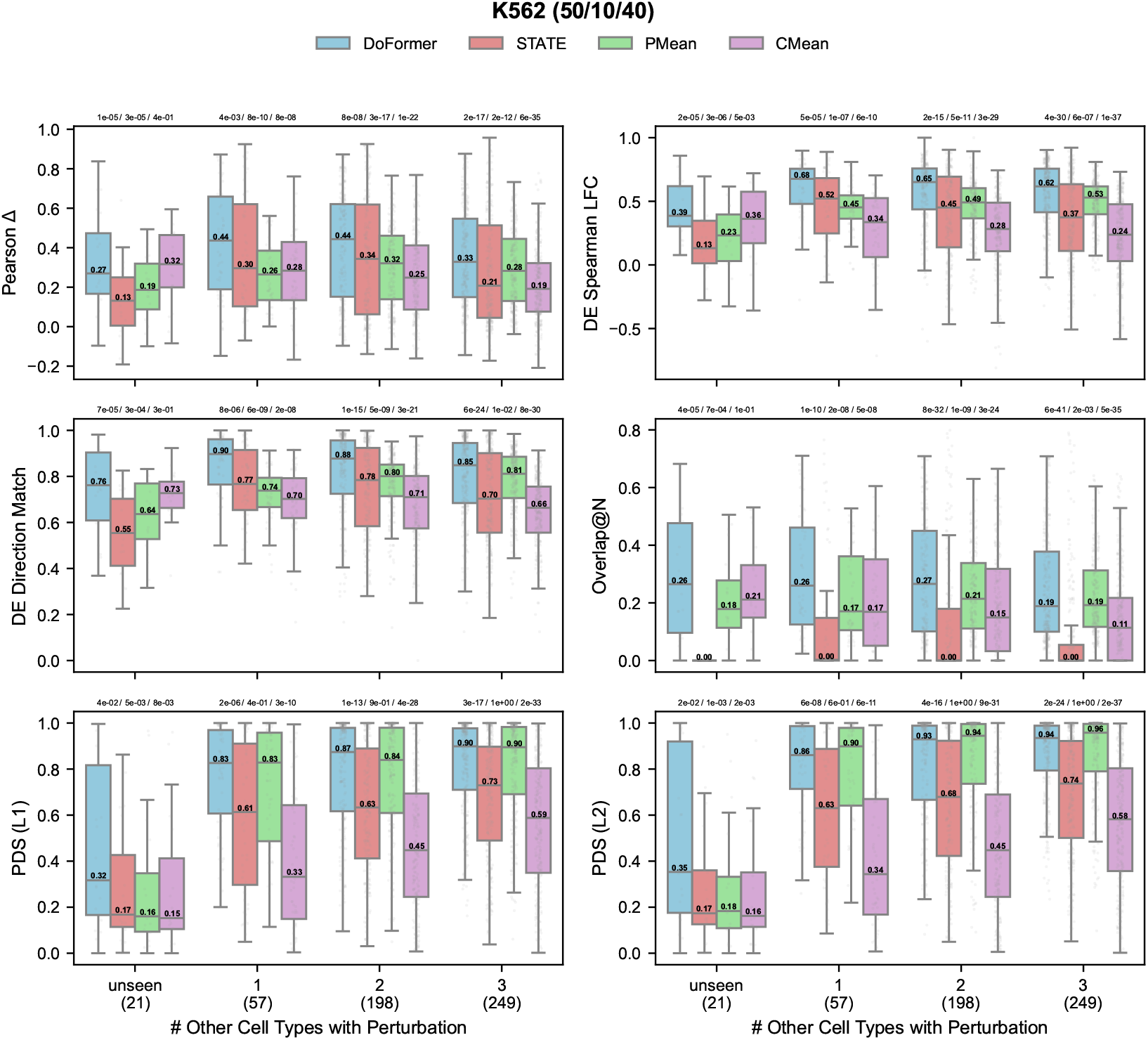
DoFormer outperforms STATE regardless of perturbations seen or unseen in other cell types. The results are for K562 in the split 50*/*10*/*40. The number of test perturbations is shown below each category (unseen, and seen in 1, 2, 3 cell types). Each point represents a perturbation, and the median of each distribution is printed. P-values from Wilcoxon signed-rank tests are shown above each distribution.

**Figure 18:**
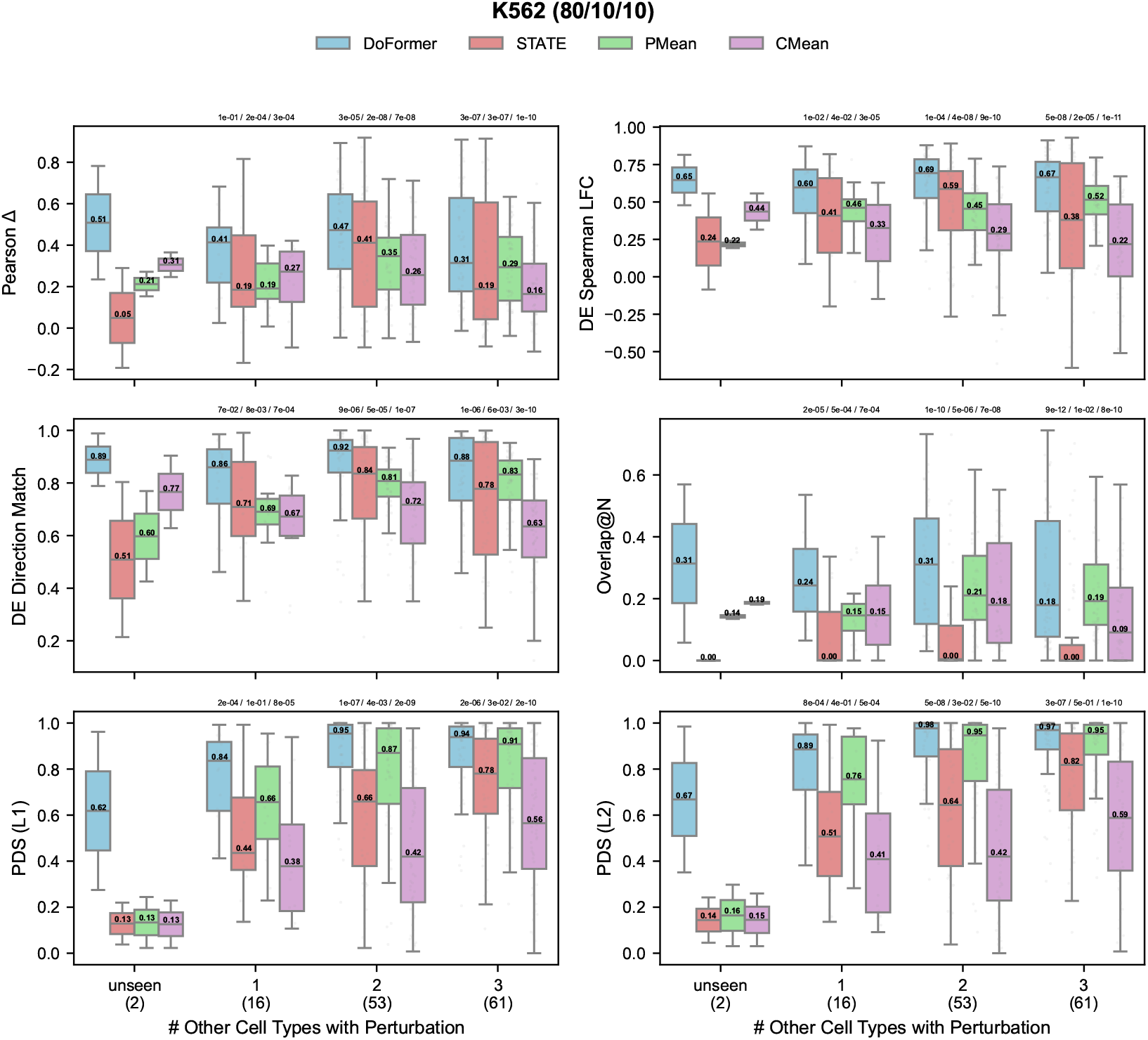
DoFormer outperforms STATE regardless of perturbations seen or unseen in other cell types. The results are for K562 in the split 80*/*10*/*10. The number of test perturbations is shown below each category (unseen, and seen in 1, 2, 3 cell types). Each point represents a perturbation, and the median of each distribution is printed. P-values from Wilcoxon signed-rank tests are shown above each distribution.

**Figure 19:**
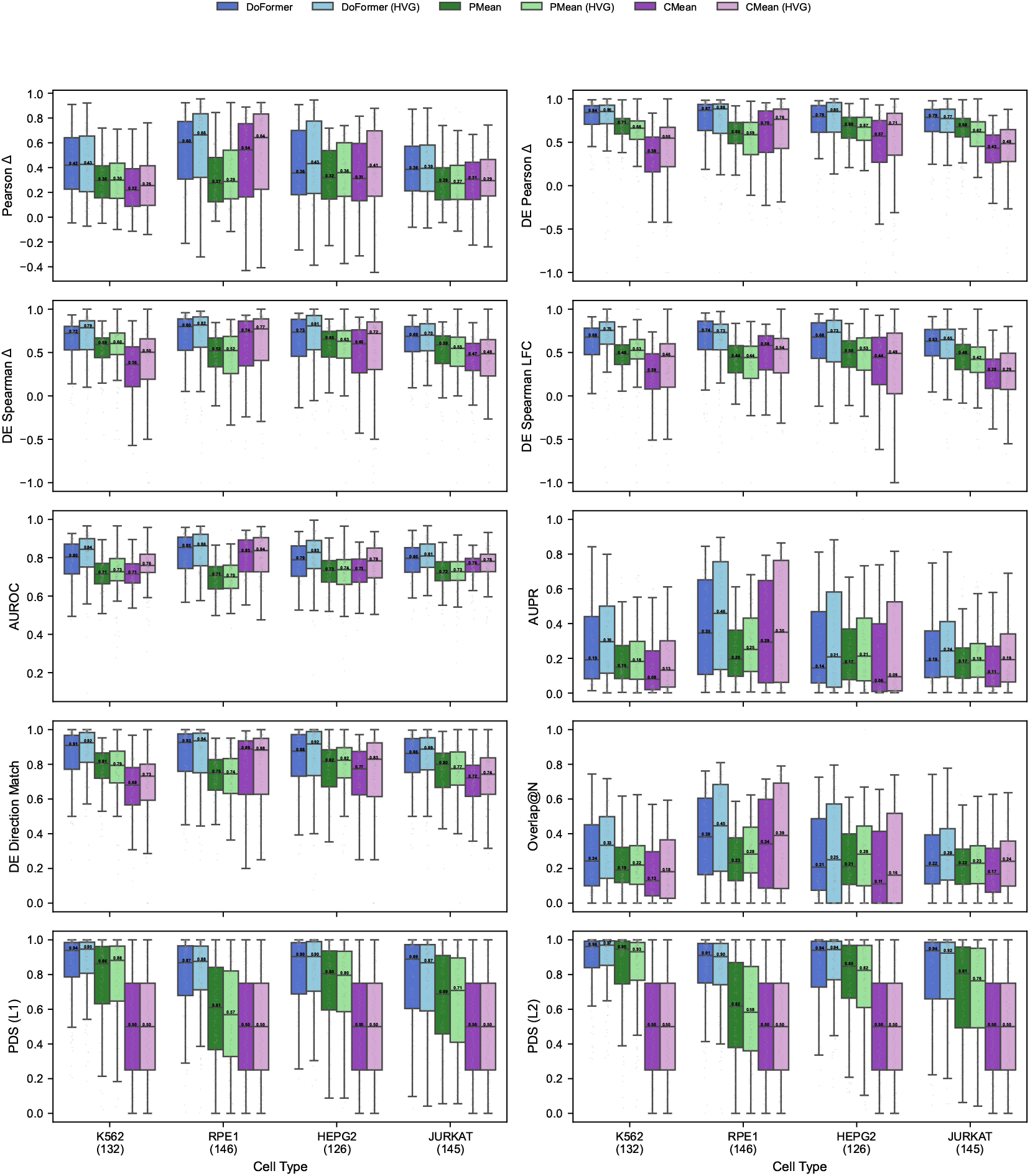
DoFormer’s evaluation metrics are better in HVGs than all genes. The results are for the split 80*/*10*/*10.

**Figure 20:**
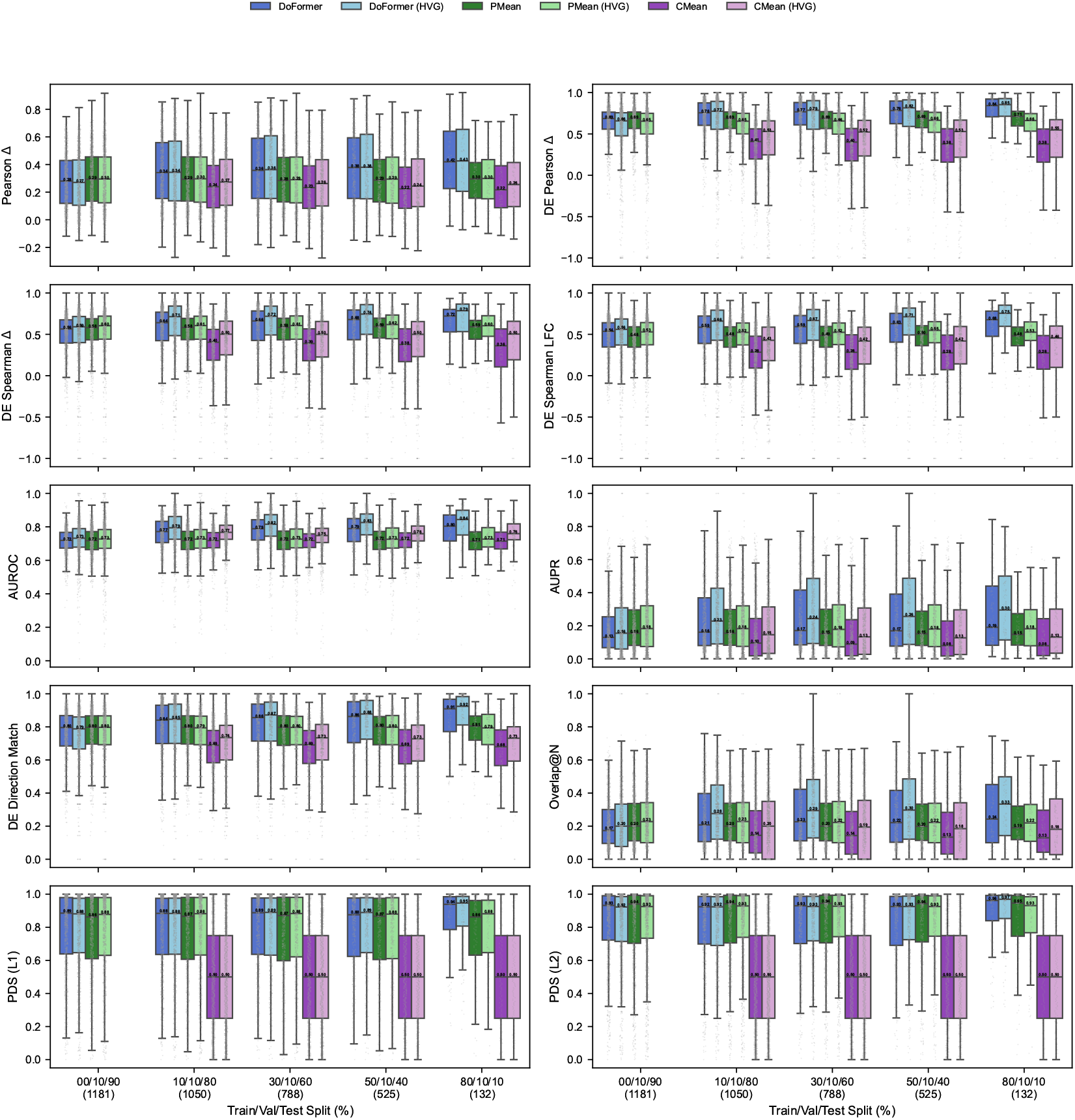
DoFormer’s evaluation metrics are better in HVGs than all genes. The results are for all the splits in K562 cells.

**Figure 21:**
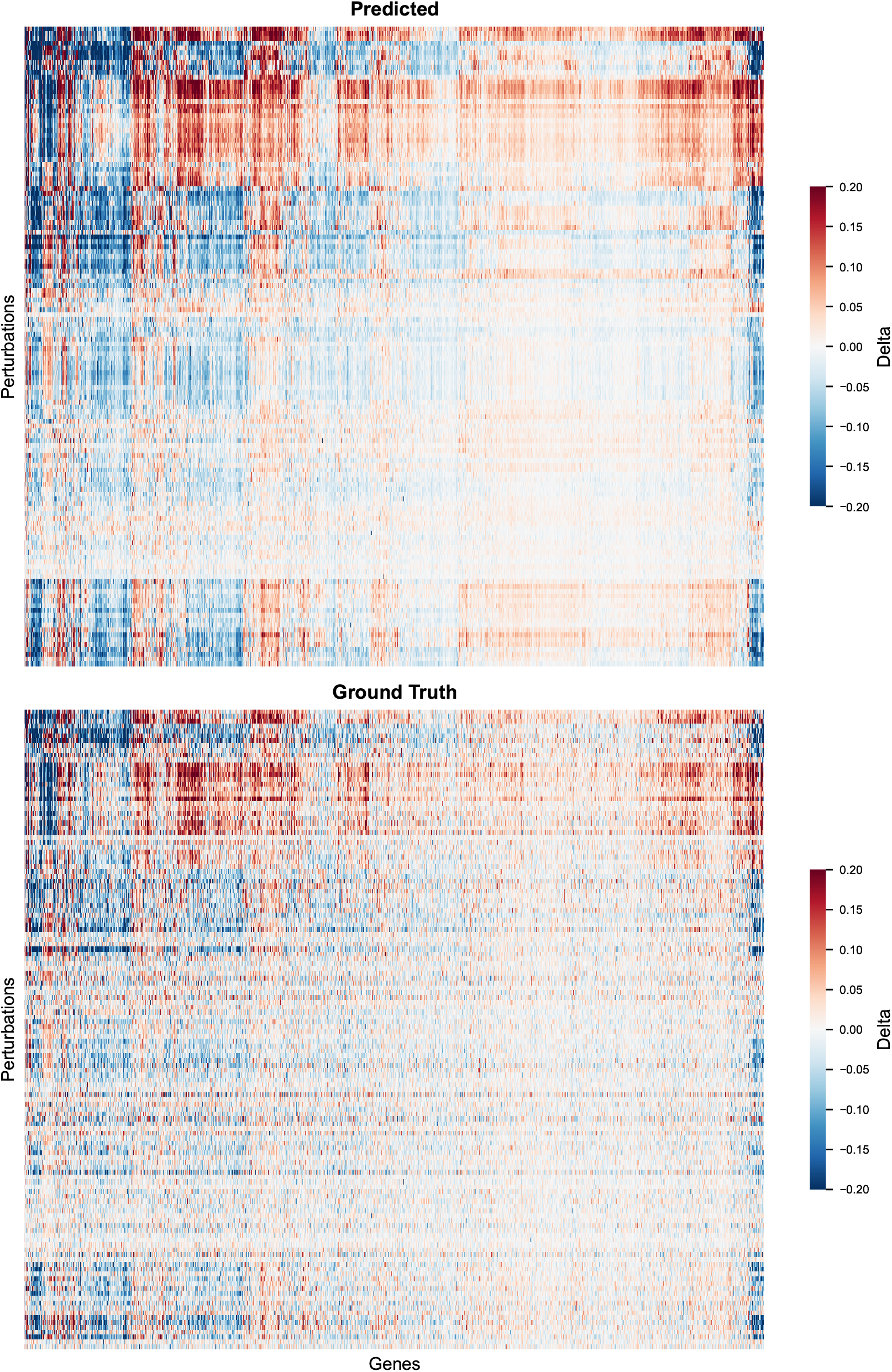
Clustermaps of DoFormer predicted and true delta values showing similarity of perturbation-gene structures, especially for the stronger effects. The results are for K562 and split 80*/*10*/*10.

**Figure 22:**
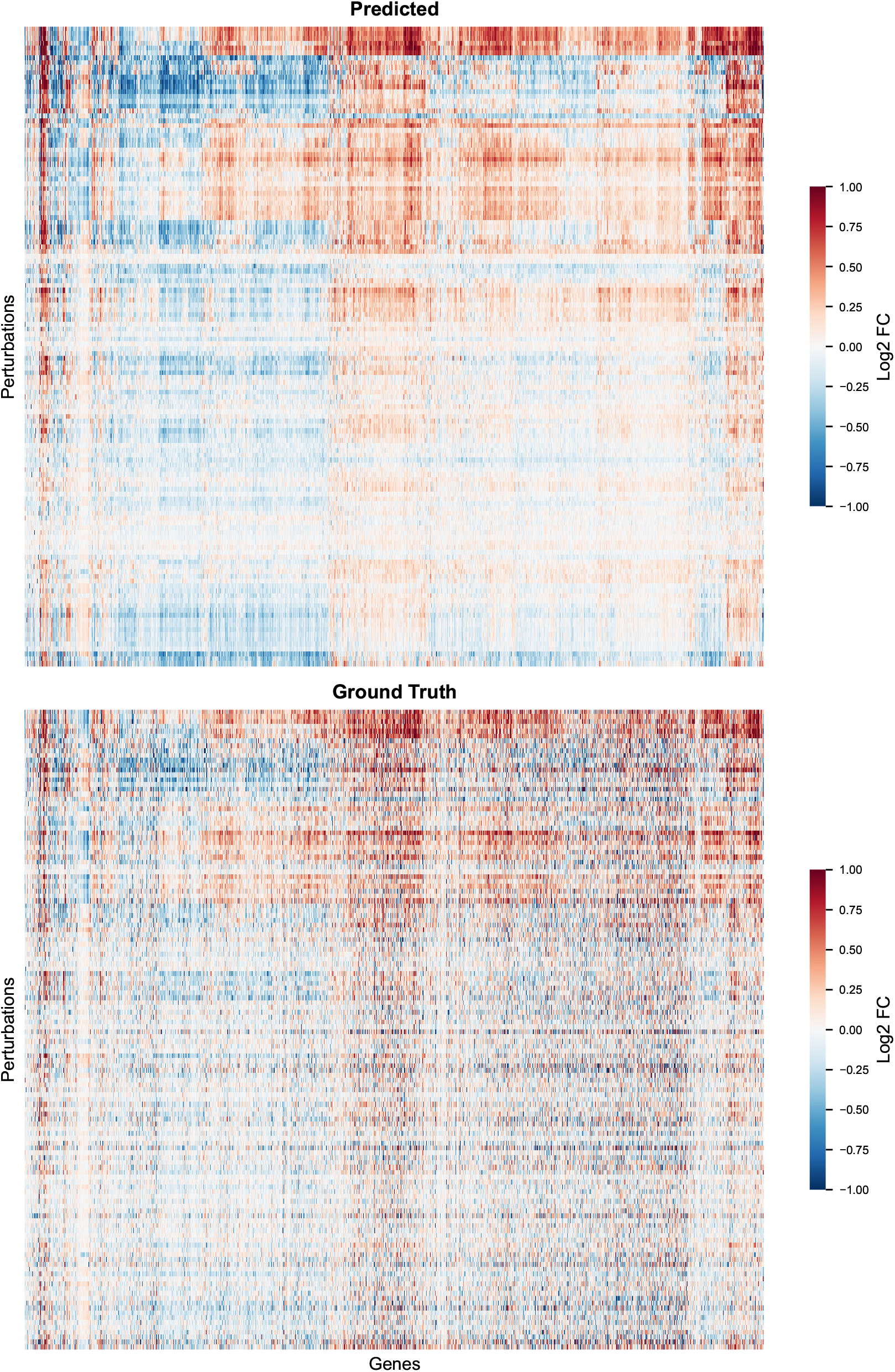
Clustermaps of DoFormer predicted and true LFC values showing similarity of perturbation-gene structures, especially for the stronger effects. The results are for K562 and split 80*/*10*/*10.

**Figure 23:**
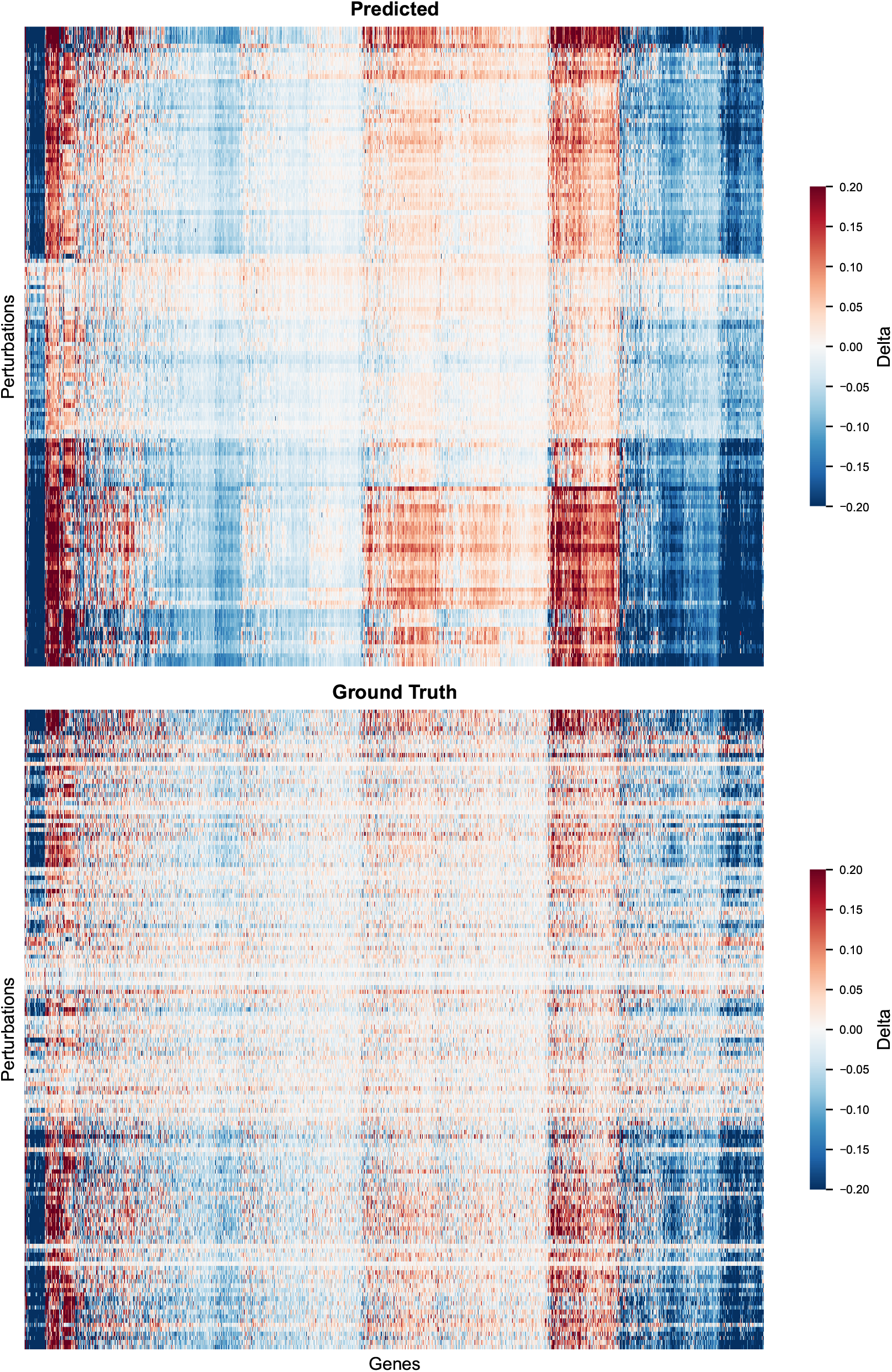
Clustermaps of DoFormer predicted and true delta values showing similarity of perturbation-gene structures, especially for the stronger effects. The results are for RPE1 and split 80*/*10*/*10.

**Figure 24:**
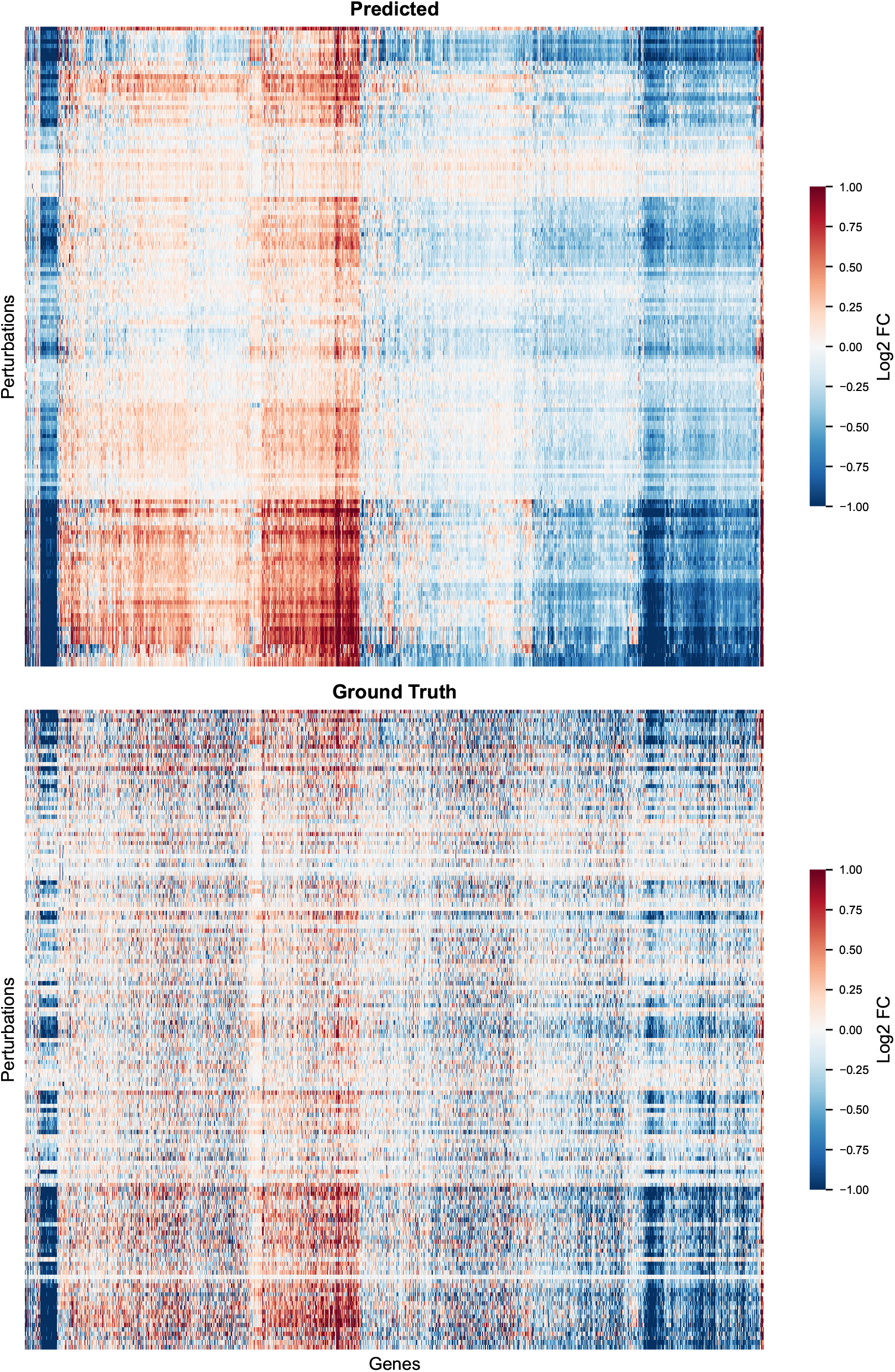
Clustermaps of DoFormer predicted and true LFC values showing similarity of perturbation-gene structures, especially for the stronger effects. The results are for RPE1 and split 80*/*10*/*10.

**Figure 25:**
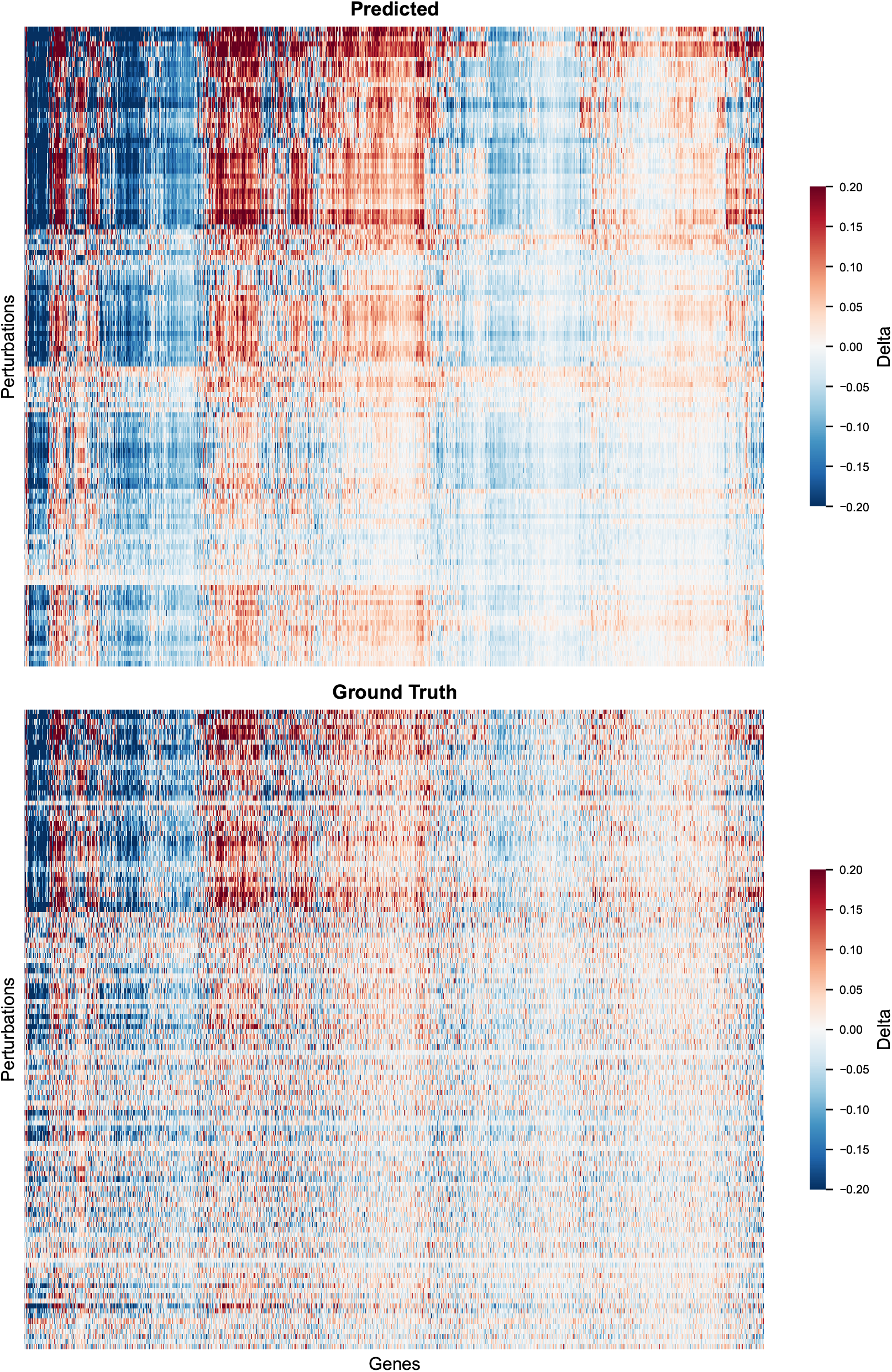
Clustermaps of DoFormer predicted and true delta values showing similarity of perturbation-gene structures, especially for the stronger effects. The results are for HEPG2 and split 80*/*10*/*10.

**Figure 26:**
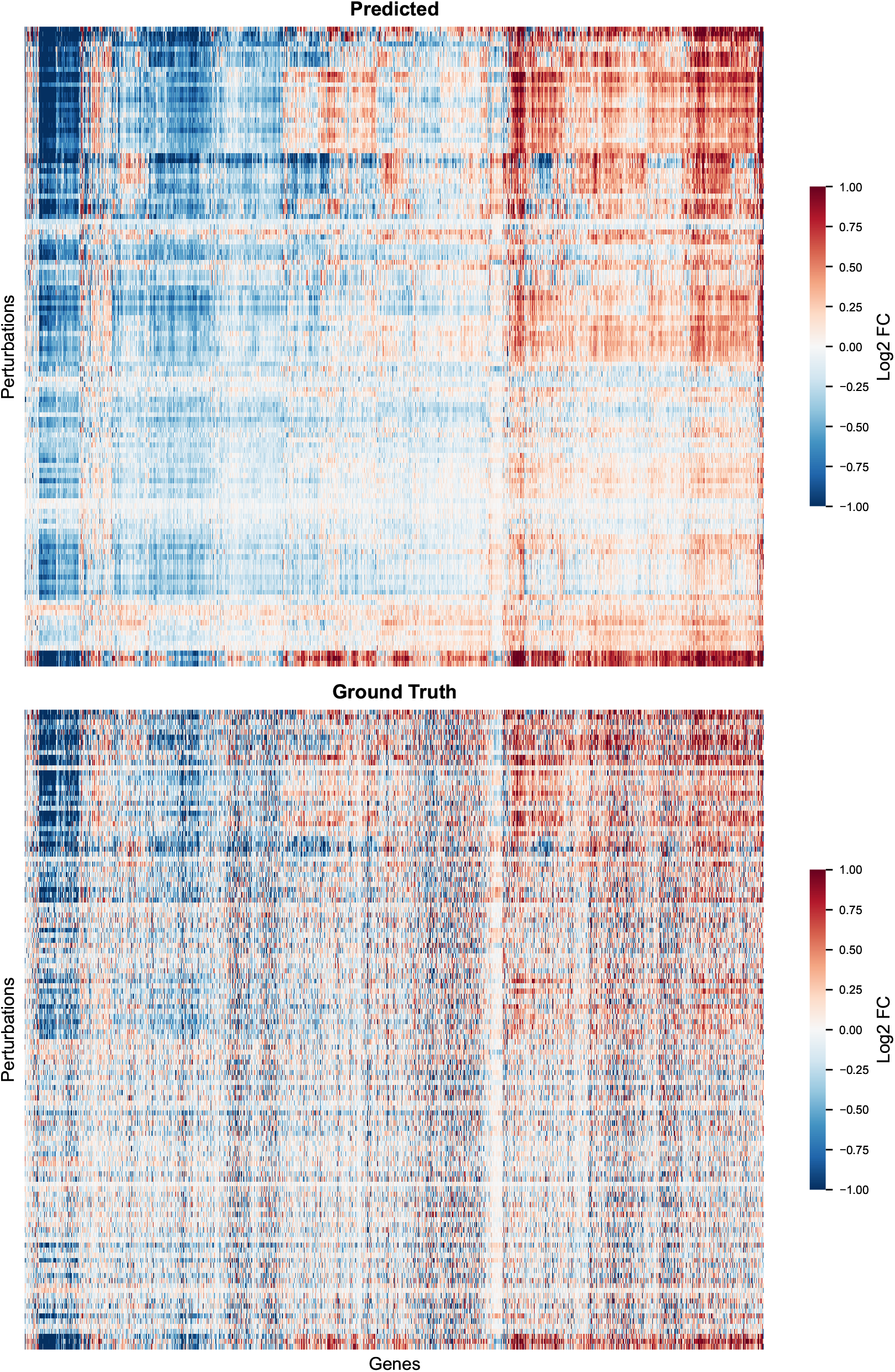
Clustermaps of DoFormer predicted and true LFC values showing similarity of perturbation-gene structures, especially for the stronger effects. The results are for HEPG2 and split 80*/*10*/*10.

**Figure 27:**
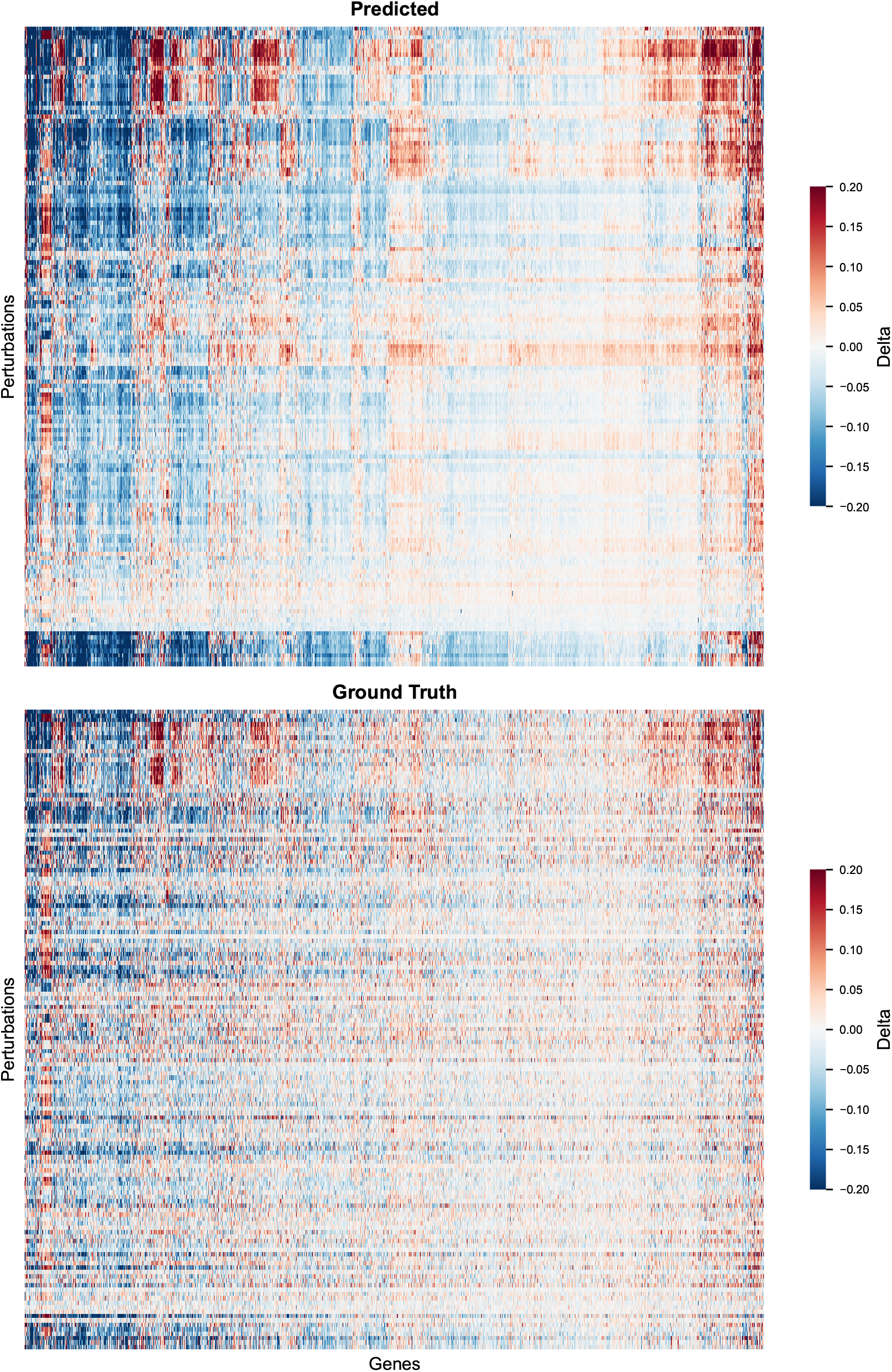
Clustermaps of DoFormer predicted and true delta values showing similarity of perturbation-gene structures, especially for the stronger effects. The results are for JURKAT and split 80*/*10*/*10.

**Figure 28:**
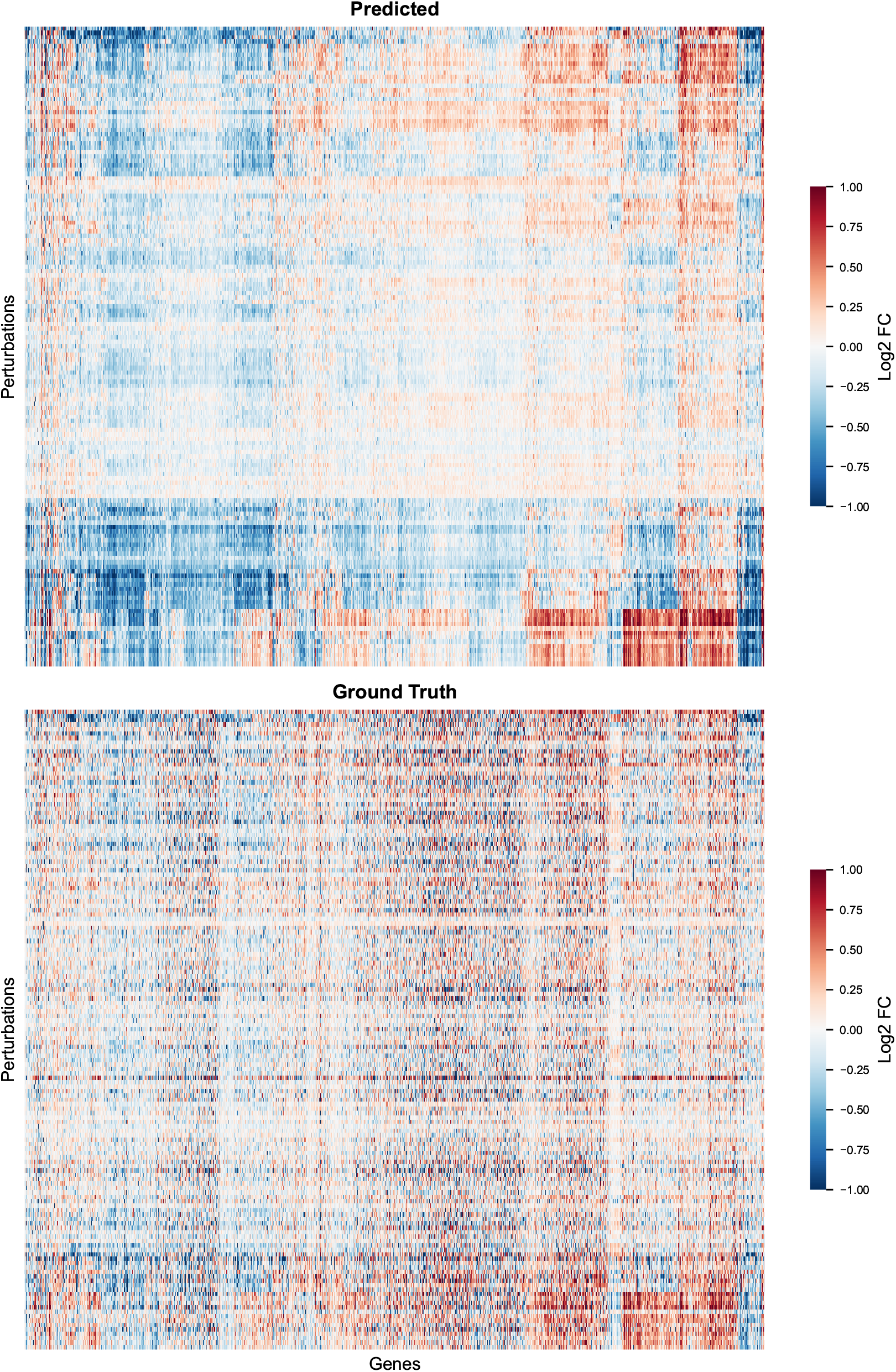
Clustermaps of DoFormer predicted and true LFC values showing similarity of perturbation-gene structures, especially for the stronger effects. The results are for JURKAT and split 80*/*10*/*10.

**Figure 29:**
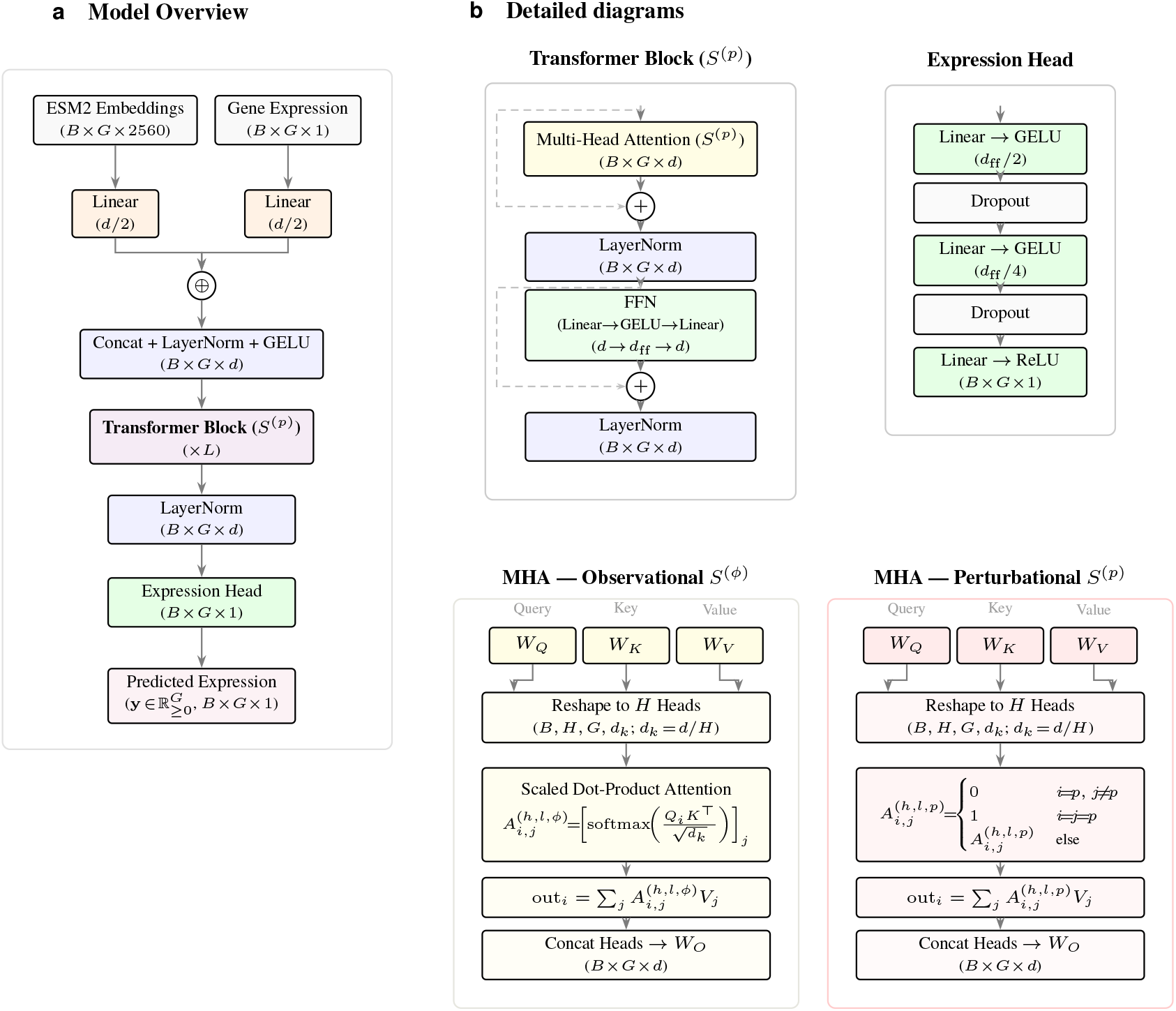
DoFormer Architecture. **(a)** Each gene is tokenized by concatenating its projected ESM embedding (*d/*2) with its projected expression value (*d/*2), normalized, and processed by *L* Transformer blocks. **(b)** The *Transformer block* applies multi-head attention with residual connections and LayerNorm. The *expression head* is a three-layer MLP with a final ReLU. In *S*^(*ϕ*)^, standard attention is used. In *S*^(*p*)^, the *do*-operator sets gene *p*’s input to *τ* and enforces 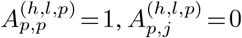 for *j*≠ *p*.

